# Distinct higher-order representations of natural sounds in human and ferret auditory cortex

**DOI:** 10.1101/2020.09.30.321695

**Authors:** Agnès Landemard, Célian Bimbard, Charlie Demené, Shihab Shamma, Sam Norman-Haignere, Yves Boubenec

## Abstract

Little is known about how neural representations of natural sounds differ across species. For example, speech and music play a unique role in human hearing, yet it is unclear how auditory representations of speech and music differ between humans and other animals. Using functional Ultrasound imaging, we measured responses in ferrets to a set of natural and spectrotemporally-matched synthetic sounds previously tested in humans. Ferrets showed similar lower-level frequency and modulation tuning to that observed in humans. But while humans showed prominent selectivity for natural vs. synthetic speech and music in non-primary regions, ferret responses to natural and synthetic sounds were closely matched throughout primary and non-primary auditory cortex, even when tested with ferret vocalizations. This finding reveals that auditory representations in humans and ferrets diverge sharply at late stages of cortical processing, potentially driven by higher-order processing demands in speech and music.

## Introduction

Surprisingly little is known about how sensory representations of natural stimuli differ across species (Theunissen and Elie, 2014). This question is central to understanding how evolution and development shape sensory representations (Moore and Woolley, 2019) as well as developing animal models of human brain functions. Audition provides a natural test case because speech and music play a unique role in human hearing (Zatorre et al., 2002; Hickok and Poeppel, 2007; Patel, 2012). While human knowledge of speech and music clearly differs from other species (Pinker and Jackendoff, 2005), it remains unclear how neural representations of speech and music differ from those in other species, particularly within the auditory cortex. Few studies have directly compared neural responses to natural sounds between humans and other animals, and those which have done so, have often observed similar responses. For example, both humans and non-human primates show regions that respond preferentially to conspecific vocalizations (Belin et al., 2000; Petkov et al., 2008). Human auditory cortex exhibits selectivity for speech phonemes (Mesgarani et al., 2014; Di Liberto et al., 2015), but much of this selectivity can be predicted by simple forms of spectrotemporal modulation tuning (Mesgarani et al., 2014), and perhaps as a consequence, can be observed in other animals such as ferrets (Mesgarani et al., 2008; Steinschneider et al., 2013). Consistent with this finding, maps of spectrotemporal modulation, measured using natural sounds, appear coarsely similar between humans and macaques (Erb et al., 2019) although temporal modulations present in speech may be over-represented in humans. Thus, it remains unclear if the representation of natural sounds in auditory cortex differs substantially between humans and other animals, and if so, how.

A key challenge is that representations of natural stimuli are transformed across different stages of sensory processing, and species may share some but not all representational stages. Moreover, responses at different sensory stages are often correlated across natural stimuli (de Heer et al., 2017), making them difficult to disentangle. Speech and music, for example, have distinctive patterns of spectrotemporal modulation energy (Singh and Theunissen, 2003; Ding et al., 2017), as well as higher-order structure (e.g. syllabic and harmonic structure) that is not well captured by modulation (Norman-Haignere and McDermott, 2018). To isolate neural selectivity for higher-order structure, we recently developed a method for synthesizing sounds whose spectrotemporal modulation statistics are closely matched to a corresponding set of natural sounds (Norman-Haignere and McDermott, 2018). Because the synthetic sounds are otherwise unconstrained, they lack perceptually salient higher-order structure, which is particularly true for complex natural sounds like speech and music which are poorly captured by modulation statistics, unlike many other natural sounds (McDermott and Simoncelli, 2011). We found that human primary auditory cortex responds similarly to natural and spectrotemporally synthetic sounds, while non-primary regions respond selectively to the natural sounds. Most of this selectivity is driven by preferential responses to natural vs. synthetic speech and music in non-primary auditory cortex. The specificity for speech and music could be due to their ecological relevance in humans and/or the fact that speech and music are more complex than other sounds, and thus perceptually differ more from their synthetic counterparts. But notably, the response preference for natural speech and music cannot be explained by speech semantics, since similar responses are observed for native and foreign speech (Norman-Haignere et al., 2015; Overath et al., 2015), or explicit musical training, since music selectivity is robust in humans without any training (Boebinger et al., 2020). These findings suggest that human non-primary regions respond selectively to higher-order acoustic features that both cannot be explained by lower-level modulation statistics and do not yet reflect explicit semantic knowledge.

The goal of the present study was to test whether such higher-order selectivity is present in other species. We test three key hypotheses: (1) higher-order selectivity in humans reflects a generic mechanism present across species for analyzing complex sounds like speech and music (2) higher-order selectivity reflects an adaptation to ecologically relevant sounds such as speech and music in humans or vocalizations in other species (3) higher-order selectivity reflects a specific adaptation in humans, potentially driven by the unique demands of speech and music perception, that is not generically present in other species even for ecologically relevant sounds. We addressed this question by measuring cortical responses in ferrets – one of the most common animal models used to study auditory cortex (Nelken et al., 2008) – to the same set of natural and synthetic sounds previously tested in humans, as well as natural and synthetic ferret vocalizations. Responses were measured using functional UltraSound imaging (fUS) (Macé et al., 2011; Bimbard et al., 2018), a newly developed wide-field imaging technique that like fMRI detects changes in neural activity via changes in blood-flow (movement of blood induces a doppler effect detectable with ultrasound). fUS has substantially better spatial resolution than fMRI making it applicable to small animals like ferrets. We found that tuning for spectrotemporal modulations present in both natural and synthetic sounds was similar between humans and animals, and could be quantitatively predicted across species, consistent with prior findings (Mesgarani et al., 2008; Erb et al., 2019). But unlike humans, ferret responses to natural and synthetic sounds were similar throughout primary and non-primary auditory cortex even when comparing natural and synthetic ferret vocalizations; and the small differences that were present in ferrets were weak and spatially scattered, unlike the selectivity observed in humans. This finding reveals that auditory representations in humans and ferrets diverge substantially at late stages of acoustic processing.

## Results

### Experiment I: Comparing ferret cortical responses to natural versus synthetic sounds

We measured cortical responses with fUS to the same 36 natural sounds tested previously in humans plus 4 additional ferret vocalizations (Experiment II tested many more ferret vocalizations). The 36 natural sounds included speech, music, and other environmental sounds (see **Table S1**). For each natural sound, we synthesized 4 sounds that were matched on acoustic statistics of increasing complexity (**Fig 1A**): (1) cochlear energy statistics (2) temporal modulation statistics (3) spectral modulation statistics and (4) spectrotemporal modulation statistics. Cochlear-matched sounds had a similar frequency spectrum, but their modulation content was unconstrained and thus differed from the natural sounds. Modulation-matched sounds were additionally constrained in their temporal and/or spectral modulation rates, measured by linearly filtering a cochleagram representation with filters tuned to different modulation rates (modulation-matched sounds also had matched cochlear statistics in order to isolate the contribution of modulation). The modulation-matched sounds audibly differ from their natural counterparts, particularly for complex sounds like speech and music that contain higher-order structure not captured by frequency and modulation statistics (listen to example sounds [here](http://mcdermottlab.mit.edu/svnh/model-matching/Stimuli_from_Model-Matching_Experiment.html)). We focused on time-averaged statistics because the hemodynamic response measured by both fMRI and fUS reflects a time-averaged measure of neural activity. As a consequence, each of the synthetic sounds can be thought of as being matched under a different model of the fUS or fMRI response (Norman-Haignere and McDermott, 2018).

**Figure 1.**
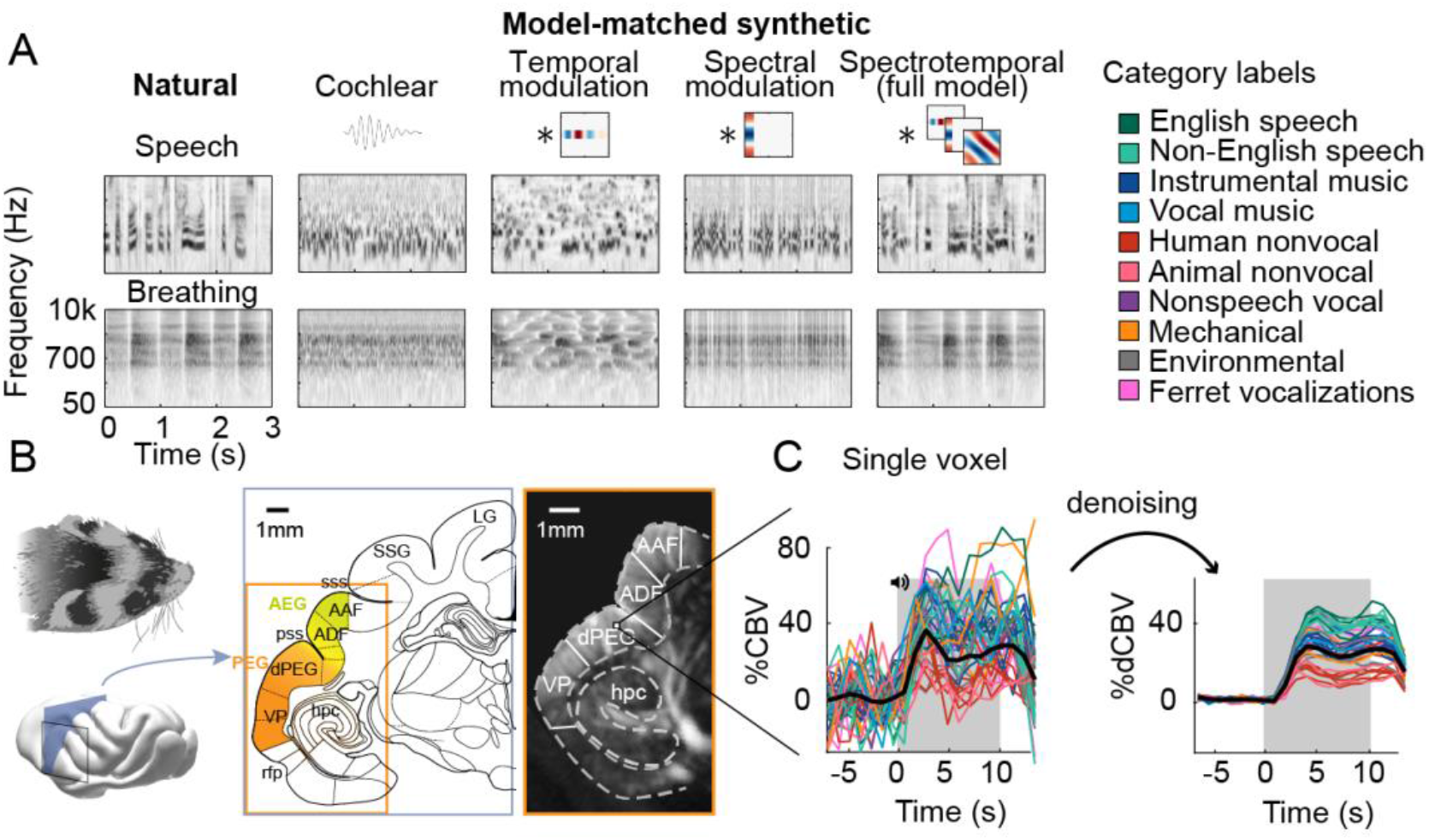
Schematic of stimuli and imaging protocol. **A**, Cochleagrams for two example natural sounds (left column) and corresponding synthetic sounds (right four columns) that were matched to the natural sounds along a set of acoustic statistics of increasing complexity. Statistics were measured by filtering a cochleagram with filters tuned to temporal, spectral or joint spectrotemporal modulations. The natural sounds were diverse, and were grouped into 10 different categories shown at right. English and Non-English speech are separated out because all of the human subjects tested in our prior study were native English speakers, and so the distinction is meaningful in humans. **B**, Schematic of the imaging procedure. A three-dimensional volume covering all of ferret auditory cortex was acquired through successive coronal slices. Auditory cortical regions (colored regions) were mapped with anatomical and functional markers. The rightmost image shows a single ultrasound image with overlaid region boundaries. Auditory regions: dPEG: dorsal posterior ectosylvian gyrus; AEG: anterior ectosylvian gyrus; VP: ventral posterior auditory field; ADF: anterior dorsal field; AAF: anterior auditory field. Non-auditory regions: hpc: hippocampus; SSG: suprasylvian gyrus; LG: lateral gyrus. Anatomical markers: pss: posterior sylvian sulcus; sss: superior sylvian sulcus. **C**, Response timecourse of a single voxel to all natural sounds, measured from raw (left) and denoised data (right). Each line reflects a different sound, and its color indicates the sound’s category. The gray region shows the time window when sound was present. The location of this voxel corresponds to the highlighted voxel in panel B.

We measured fUS responses throughout primary and non-primary ferret auditory cortex (**Fig 1B**). We first plot the response timecourse to all 40 natural sounds for one example voxel in non-primary auditory cortex (dPEG) (**Fig 1C**). We plot the original timecourse of the voxel as well as a denoised version computed by projecting the timecourse onto a small number of reliable components, which we found substantially improved prediction accuracy in left-out data (see Methods for details). As expected and similar to fMRI, we observed a gradual build-up of the hemodynamic response after stimulus onset. The shape of the response timecourse was similar across stimuli, but the magnitude of the response varied, and we thus summarized the response of each voxel to each sound by its time-averaged response magnitude (the same approach used in our prior fMRI study).

We next plot the time-averaged response of two example voxels – one in primary auditory cortex (A1) and one in a non-primary area (dPEG) – to natural and corresponding synthetic sounds that have been matched on the full spectrotemporal modulation model (**Fig 2A**). For comparison, we plot the test-retest reliability of each voxel across repeated presentations of the same sound (**Fig 2B**), as well as corresponding figures from two example voxels in human primary/non-primary auditory cortex (**Fig 2C-D**; these voxels are re-plotted from our prior paper). As in our prior study, we quantified the similarity of responses to natural and synthetic sounds using the normalized squared error (NSE). The NSE takes a value of 0 if responses to natural and synthetic sounds are the same, and 1 if there is no correspondence between the two (see Methods for details).

**Figure 2:**
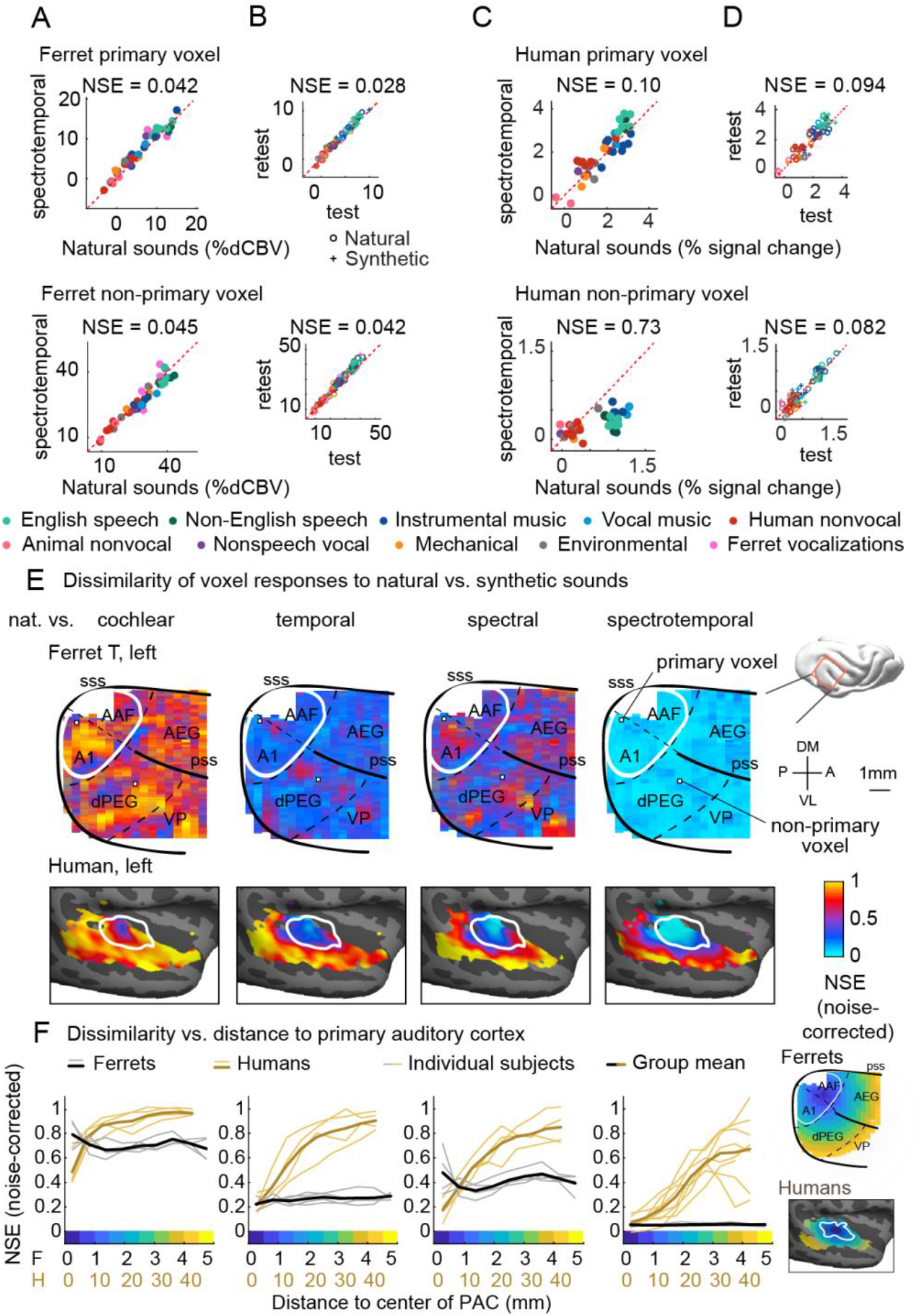
Dissimilarity of responses to natural vs. synthetic sounds in ferrets and humans. **A**, Response of two example fUS voxels to natural and corresponding synthetic sounds with matched spectrotemporal modulation statistics. Each dot shows the time-averaged response to a single pair of natural/synthetic sounds (after denoising), with colors indicating the sound category. The example voxels come from primary (top, A1) and non-primary (bottom, dPEG) regions of the ferret auditory cortex. The normalized squared error (NSE) quantifies the dissimilarity of responses. **B**, Test-retest response of the example voxels across all natural (o) and synthetic (+) sounds (odd vs. even repetitions). The responses were highly reliable due to the denoising procedure. **C-D**, Same as panel A-B, but showing two example voxels from human primary/non-primary auditory cortex. **E**, Maps plotting the dissimilarity of responses to natural vs. synthetic sounds from one ferret hemisphere (top row) and from humans (bottom row). Each column shows results for a different set of synthetic sounds. The synthetic sounds were constrained by statistics of increasing complexity from left to right: just cochlear statistics, cochlear + temporal modulation statistics, cochlear + spectral modulation statistics, and cochlear + spectrotemporal modulation statistics. Dissimilarity was quantified using the normalized squared error (NSE), corrected for noise using the test-retest reliability of the voxel responses. Ferret maps show a “surface” view from above of the sylvian gyri, similar to the map in humans. Surface views were computed by averaging activity perpendicular to the cortical surface. The border between primary and non-primary auditory cortex is shown with a white line in both species, and was defined using tonotopic gradients. Areal boundaries in the ferret are also shown (dashed thin lines). This panel shows results from one hemisphere of one animal (Ferret T, left hemisphere), but results were similar in other animals/hemispheres (**Fig S1**). The human map is a group map averaged across many subjects, but results were similar in individual subjects (Norman-Haignere and McDermott, 2018). **F**, Voxels were binned based on their distance to primary auditory cortex (defined tonotopically). This figure plots the median NSE value in each bin. Each thin line corresponds to a single ferret hemisphere (gray) or a single human subject averaged across hemispheres (gold) (results were very similar in the left and right hemisphere of humans). Thick lines show the average across all hemispheres/subjects.

Both the primary and non-primary ferret voxels produced nearly identical responses to natural and corresponding synthetic sounds (NSEs: 0.042, 0.045), suggesting that spectrotemporal modulation are sufficient to account for the responses in these voxels. The human primary voxel also showed similar responses to natural and synthetic responses, and the NSE for natural vs. synthetic sounds (0.1) was similar to the test-retest NSE (0.094), indicating that the response was about as similar as possible given the noise ceiling. In contrast, the human non-primary voxel responded substantially more to the natural speech (green) and music (blue) than matched synthetics, yielding a high NSE value (0.73). This pattern demonstrates that spectrotemporal modulations are insufficient to drive the response of the human non-primary voxel, plausibly because it responds to higher-order features that are not captured by modulation statistics.

We quantified this trend across voxels by plotting maps of the noise-corrected NSE between natural and synthetic sounds (**Fig 2E** shows one hemisphere of one animal, but results were very similar in other hemispheres of other animals, see **Fig S1**). We show separate maps for each of the different sets of statistics used to constrain the synthetic sounds (cochlear, temporal modulation, spectral modulation and spectrotemporal modulation). Each map shows a view from above auditory cortex, computed by averaging NSE values perpendicular to the cortical sheet. We summarized the data in this way, because we found that maps were very similar across the different layers within a cortical column. Below we plot corresponding maps from humans. The human maps are based on data averaged across subjects, but similar results were observed in individual subjects (Norman-Haignere and McDermott, 2018).

In ferrets, we observed a similar pattern throughout both primary and non-primary regions: responses became more similar as we matched additional acoustic features with NSE values close to 0 for sounds matched on the full spectrotemporal model. This pattern contrasts sharply with that observed in humans, where we observed a clear and substantial rise in NSE values when moving from primary to non-primary auditory cortex even for sounds matched on joint spectrotemporal modulations statistics. We quantified these effects by measuring NSE values using ROIs binned based on distance to primary auditory cortex, as was done previously in humans (**Fig 2F**). This analysis revealed a substantial and significant rise in NSEs when matching additional acoustic features in ferrets (NSE spectrotemporal < NSE temporal < NSE spectral < NSE cochlear, p < 0.01 via a bootstrapping analysis across the sound set). But there was little difference in NSEs between ferret primary and non-primary regions, with NSE values close to zero in all regions for spectrotemporally matched synthetics. In contrast, every human subject tested showed larger NSE values in non-primary regions, yielding a significant species difference (p < 0.01 via a sign-test comparing each ferret to all of the human subjects tested; see Methods for details). This finding demonstrates that higher-order selectivity for complex natural sounds like speech and music is not a generic feature of higher-order processing in mammals.

### Assessing and comparing selectivity for frequency and modulation across species

Our NSE maps suggest that ferret cortical responses are selective for frequency and modulation, but do not reveal how this selectivity is organized or whether it is similar to that in humans. While it is not feasible to inspect or plot all individual voxels, we found that fUS responses like human fMRI responses are low-dimensional and can be explained as the weighted sum of a small number of component response patterns. This observation served as the basis for our denoising procedure, as well as a useful way to examining ferret cortical selectivity and comparing that selectivity with humans. We found that we could discriminate approximately 8 distinct component response patterns before over-fitting to noise (**Fig S2C**).

We first examined the selectivity of the inferred response patterns and their anatomical distribution of weights in the brain (**Fig 3** shows three example components; **Fig S3** shows all 8 components). All of the component response profiles showed significant correlations with measures of energy at different cochlear frequencies and spectrotemporal modulation rates (**Fig 3D-E**) (p < 0.01 for all components for both frequency and modulation features; statistics computed via a permutation test across the sound set). Two components (f1 & f2) had responses that correlated with energy at high and low-frequencies respectively, with voxel weights that mirrored the tonotopic gradients measured in these animals (compare **Fig 3B** and **3A;** see **Fig S4** for all hemispheres/animals), similar to the tonotopic components previously identified in humans (Norman-Haignere et al., 2015) (**Fig S5**, components h1 and h2). We also observed components with weak frequency tuning but prominent tuning for spectrotemporal modulations (**Fig S3**), again similar to humans. Perhaps surprisingly, one component (f3) responded selectively to speech sounds, and its response correlated with energy at frequency and modulation rates characteristic of speech (insets in **Fig 3D-E**, bottom row). But notably, all of the inferred components, including the speech-selective component, produced very similar responses to natural and synthetic sounds (**Fig 3C**), suggesting that their selectivity can be explained by their tuning for frequency and modulation. This contrasts with the speech- and music-selective components previously observed in humans, which responded selectively to natural speech and music, respectively, and which clustered in distinct non-primary regions of human auditory cortex (see **Fig S5**, components h5 and h6). This finding shows that selectivity for natural speech compared with other natural sounds is in fact not unique to humans, and thus that comparing responses to natural vs. synthetic sounds is critical to revealing representational differences between species.

**Figure 3:**
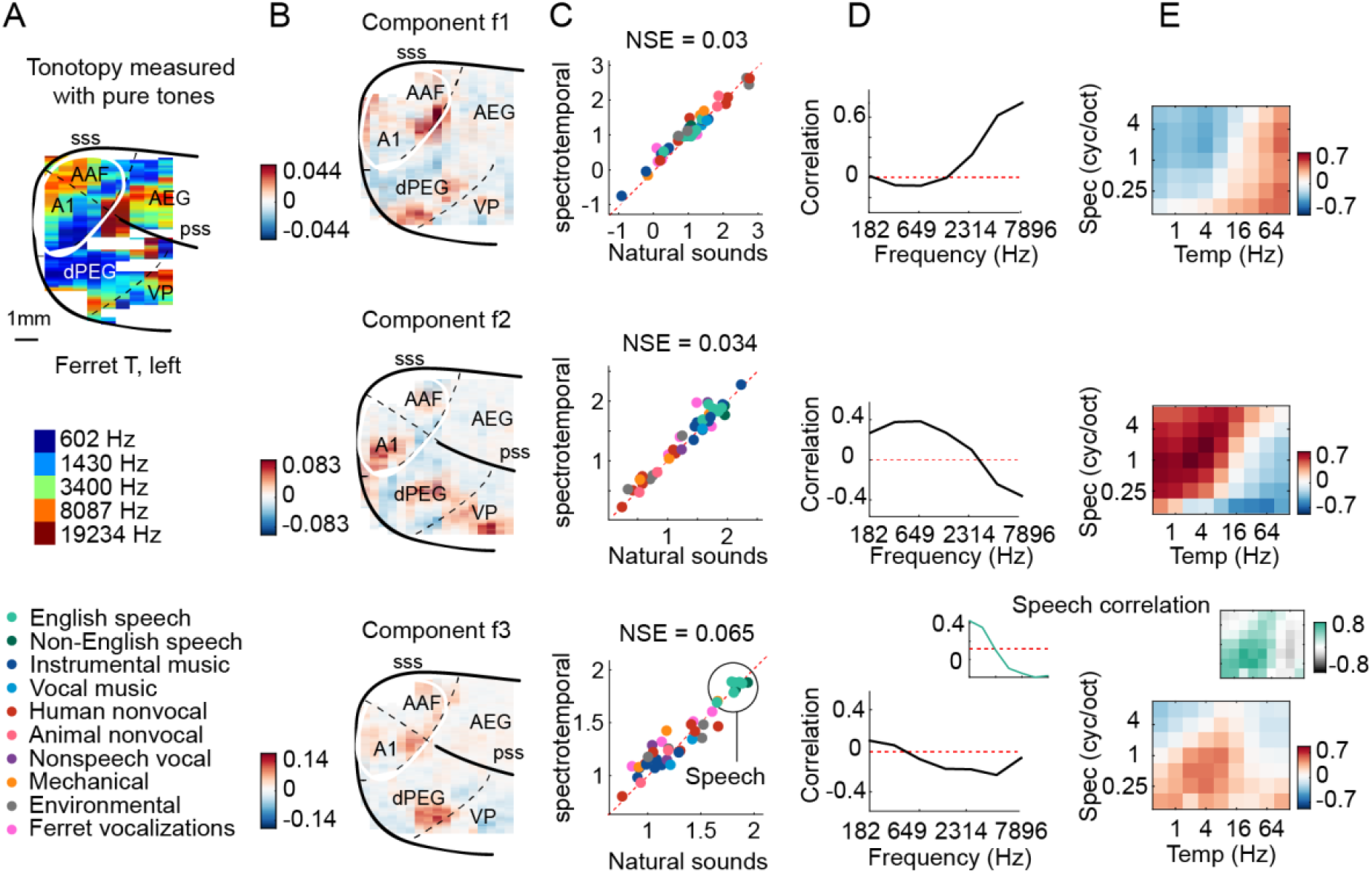
Organization of frequency and modulation selectivity in ferret auditory cortex, revealed by component analysis. **A**, For reference with the weight maps in panel B, a tonotopic map is shown, measured using pure tones. The map is from one hemisphere of one animal (Ferret T, left). **B**, Voxel weight maps from three components, inferred using responses to natural and synthetic sounds (see **Fig S3** for all 8 components and **Fig S4** for all hemispheres). Each map was computed by averaging weights perpendicular to the cortical surface, which was done because the weights were very similar across layers within a column (see **Fig S4C**). The maps for components f1 and f2 closely mirrored the high and low-frequency tonotopic gradients respectively. **C**, Component response to natural and spectrotemporally-matched synthetic sounds, colored based on category labels (labels shown at the bottom left of the figure). Components f1 and f2 did not respond selectively to particular categories. Component f3 responded preferentially to speech sounds. **D**, Correlation of component responses with energy at different audio frequencies, measured from a cochleagram. Inset for f3 shows the correlation pattern that would be expected from a response that was perfectly selective for speech (i.e. 1 for speech, 0 for all other sounds). **E**, Correlations with modulation energy at different temporal and spectral rates. Inset shows the correlation pattern that would be expected for a perfectly speech-selective response.

Overall, the frequency and modulation selectivity evident in the ferret components appeared similar to that in humans (Norman-Haignere et al., 2015). To quantitatively evaluate similarity, we attempted to predict the response of each human component, inferred from our prior work, from those in the ferrets (**Fig S6**) and vice versa (**Fig S7**). We found that much of the component response variation to synthetic sounds could be predicted across species (**Fig S6B&D, S7A&C**). This finding is consistent with the hypothesis that tuning for frequency and modulation is similar across species, since the synthetic sounds only varied in their frequency and modulation statistics. In contrast, differences between natural vs. synthetic sounds were only robust in humans and as a consequence could not be predicted from responses in ferrets (**Fig S6C&E**). Thus, selectivity for frequency and modulation is both qualitatively and quantitatively similar across species, despite large and substantial differences in higher-order tuning.

### Experiment II: Testing the importance of ecological relevance

The results of Experiment I show that higher-order selectivity in humans is not a generic feature of auditory processing for complex sounds. However, the results could still be explained by a difference in ecological relevance, since differences between natural and synthetic sounds in humans are mostly driven by speech and music (Norman-Haignere and McDermott, 2018) and Experiment I included more speech (8) and music (10) sounds than ferret vocalizations (4). To test this possibility, we performed a second experiment that included many more ferret vocalizations (30) (**Fig 4A**), as well as a smaller number of speech (14) and music (16) sounds to allow comparison with Experiment I. We only synthesized sounds matched in their full spectrotemporal modulation statistics to be able to test a broader sound set.

**Figure 4.**
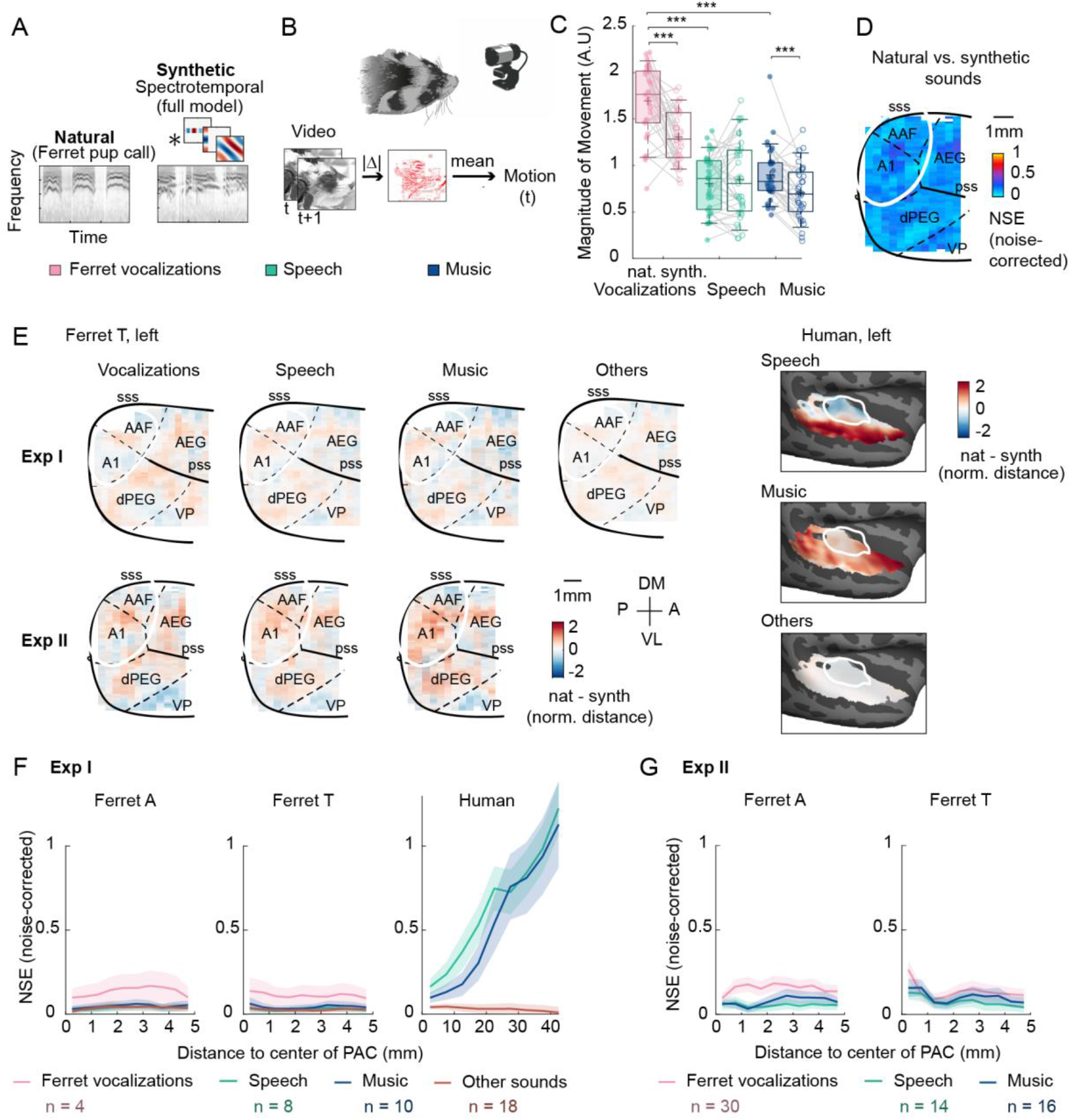
Testing the importance of ecological relevance. **A**, Experiment II measured responses to a much larger number of ferret vocalizations (30), as well as a smaller number of speech (14) and music (16) sounds, unlike Experiment I which only tested 4 ferret vocalizations. Cochleagrams for an example natural and synthetic vocalization (a “pup call”) are plotted. **B**, The animal’s spontaneous movements were monitored with a video recording of the animal’s face. Motion was measured as the mean absolute deviation between adjacent video frames, averaged across pixels. **C**, Average evoked movement amplitude for natural (shaded) and synthetic (unshaded) sounds broken down by category. Each dot represents one recording session. Significant differences between natural and synthetic sounds, and between categories of natural sounds are plotted (paired Wilcoxon test, p<0.001: ***). Evoked movement amplitude was normalized by the standard deviation across sounds for each recording session prior to averaging across sound category (necessary because absolute pixel deviations cannot be meaningfully compared across sessions). Results were consistent across ferrets (**Fig S8A**). Both animals moved substantially more during natural ferret vocalizations compared with both matched synthetics as well as speech and music. **D**, Map showing the dissimilarity between natural and spectrotemporally matched synthetic sounds from Experiment II for one hemisphere (Ferret T, left; see **Fig S8B** for all hemispheres), measured using the noise-corrected NSE across sounds. NSE values were low across auditory cortex, replicating the first experiment. **E**, Maps showing the average difference between responses to natural and synthetic sounds for vocalizations, speech, music, and others sounds, normalized for each voxel by the standard deviation across all sounds. Results are shown for the same ferret hemisphere (T, left) for both Experiment I and II. Humans were only tested in Experiment I. **F**, NSE for different sound categories, plotted as a function of distance to primary auditory cortex (binned as in **Fig 2F**). Shaded area represents +/- 1 s.e.m. (**Fig S8D** plots NSEs for individual sounds) **G**, Same as panel **F** but showing results from Experiment II.

Using a video recording of the animals’ face (**Fig 4B**), we found that the ferrets showed greater spontaneous movements during the presentation of the natural ferret vocalizations compared with both the synthetic sounds and the other natural sounds (**Fig 4C;** see **Fig S8** for additional plots from individual animals and finer-grained vocalization categories). This observation demonstrates that natural ferret vocalizations contain additional structure that is missing from their synthetic counterparts, and that this additional structure is sufficiently salient to cause a spontaneous increase in motion without any overt training. Moreover, the behavioral differences between natural and synthetic vocalizations were greater than those for speech (p < 0.001 via Wilcoxon signed-rank test) and music (p < 0.05), demonstrating that the additional structure present in vocalizations is more salient to the ferret than the additional structure present in natural speech and music. To prevent this motion from affecting the ultrasound responses, we designed a denoising procedure that greatly minimized correlations between the ultrasound responses and motion without removing sound-evoked activity (see Methods and Appendix).

Despite this clear behavioral difference, we nonetheless found that voxel responses to natural and synthetic sounds were similar throughout primary and non-primary regions, yielding small NSE values (**Fig 4D**). This result demonstrates that our key findings from Experiment I are not due to the weak ecological relevance of the tested sounds, since a qualitatively similar result was obtained in Experiment II when half of the sounds were ferret vocalizations.

To directly test if ferrets showed selective responses to natural vs. synthetic ferret vocalizations, we computed maps showing the average difference between natural vs. synthetic sounds for different categories, using data from both Experiments I and II (**Fig 4E**). We also separately measured the NSE for sounds from different categories (**Fig 4F-G**; note the normalization term in the NSE was computed using all sounds to avoid inadvertently normalizing out meaningful differences between sounds/categories). We plot the median NSE for sounds from different categories as a function of distance to primary auditory cortex for each animal and experiment (**Fig 4F-G; Fig S8D** shows the distribution of NSE values for individual sound pairs). This analysis revealed that NSE values in ferrets were slightly elevated for ferret vocalizations compared with other categories (**Fig 4F-G**), consistent with their ecological relevance. This effect, however, was small and inconsistent, reaching significance in only one of the two animals in Experiment II (Ferret A, p < 0.005, Wilcoxon test) (the effect was significant in both animals in Experiment I, but this experiment only tested 4 ferret vocalizations). Moreover, the small differences that were present between natural and synthetic sounds were spatially distributed throughout primary and non-primary regions, and very similar to those for speech, music and other natural sounds (**Fig 4E**). In contrast, humans showed large and selective responses to speech and music that were concentrated in distinct non-primary regions (lateral for speech and anterior/posterior for music) and clearly different from those for other natural sounds (**Fig 4E**). Thus, ferrets do not show any of the neural signatures of higher-order selectivity that we previously identified in humans (large effect size, spatially clustered responses, and a clear non-primary bias), even for con-specific vocalizations, which produced clear behavioral differences reflecting their ecological significance.

## Discussion

Our study reveals a prominent divergence in the representation of natural sounds between humans and ferrets. Using a recently developed wide-field imaging technique (functional Ultrasound), we measured cortical responses in the ferret to a set of natural and spectrotemporally-matched synthetic sounds previously tested in humans. We found that selectivity for frequency and modulation statistics in the synthetic sounds was similar across species. But unlike humans, who showed selective responses to natural vs. synthetic speech and music in non-primary regions, ferrets cortical responses to natural and synthetic sounds were similar throughout primary and non-primary auditory cortex, even when tested with ferret vocalizations. This finding suggests that higher-order selectivity in humans for natural vs. synthetic speech/music (1) does not reflect a species-generic mechanism for analyzing complex sounds and (2) does not reflect a species-generic adaptation for coding ecologically relevant sounds like con-specific vocalizations. Instead, our findings suggest that auditory representations in humans fundamentally diverge from ferrets at higher-order processing stages, plausibly driven by the unique demands of speech and music.

### Species differences in the representation of natural sounds

The central challenge of sensory coding is that behaviorally relevant information is often not explicit in the inputs to sensory systems. As a consequence, sensory systems transform their inputs into higher-order representations that expose behaviorally relevant properties of stimuli (DiCarlo and Cox, 2007; Mizrahi et al., 2014; Theunissen and Elie, 2014). The early stages of this transformation are thought to be conserved across many species. For example, all mammals transduce sound pressure waveforms into a frequency-specific representation of sound energy in the cochlea, although the resolution and frequency range of cochlear tuning differ across species (Bruns and Schmieszek, 1980; Köppl et al., 1993; Joris et al., 2011; Walker et al., 2019). But it has remained unclear whether representations at later stages are similarly conserved across species.

Only a few studies have attempted to compare cortical representations of natural sounds between humans and other animals, and these studies have typically found similar representations in auditory cortex. Studies of speech phonemes in ferrets (Mesgarani et al., 2008) and macaques (Steinschneider et al., 2013) have replicated many neural phenomena observed in humans (Mesgarani et al., 2014). A recent fMRI study found that maps of spectrotemporal modulation tuning, measured using natural sounds, are coarsely similar between humans and macaques, although slow temporal modulations which are prominent in speech were better decoded in humans compared with macaques (Erb et al., 2019), potentially analogous to prior findings of enhanced cochlear frequency tuning for behaviorally relevant sound frequencies (Bruns and Schmieszek, 1980; Köppl et al., 1993). Thus, prior work has revealed quantitative differences in the extent and resolution of neural tuning for different acoustic frequencies and modulation rates. But it has remained unclear whether there are qualitative differences in how natural sounds are represented across species.

Our study demonstrates that human non-primary regions exhibit a form of higher-order acoustic selectivity that is almost completely absent in ferrets. Ferret cortical responses to natural and spectrotemporally matched synthetic sounds were closely matched throughout their auditory cortex, and the small differences that we observed were scattered throughout primary and non-primary regions (**Fig 4E**), unlike the pattern observed in humans. As a consequence, the differences that we observed between natural and synthetic sounds in humans were not predictable from cortical responses in ferrets (**Fig S6C**), even though we could predict responses to synthetic sounds across species (**Fig S6B&E**). This higher-order selectivity is unlikely to be explained by explicit semantic knowledge about speech or music, since similar responses are observed for foreign speech (Norman-Haignere et al., 2015; Norman-Haignere and McDermott, 2018) and music selectivity is robust in listeners without musical training (Boebinger et al., 2020). These results suggest that humans develop or have evolved a higher-order stage of acoustic analysis, potentially specific to speech and music, that cannot be explained by standard frequency and modulation statistics and is largely absent from the ferret brain. This specificity for speech and music could be due to their acoustic complexity, their behavioral relevance to humans, or a combination of the two.

By comparison, our study suggests that there is a substantial amount of cross-species overlap in the cortical representation of frequency and modulation features. Both humans and ferrets exhibited tonotopically organized selectivity for different frequencies. Moreover, modulation selectivity accounted for a large fraction of the cortical responses (**Fig 2E**), even in primary auditory cortex, which emphasizes the importance of modulation tuning in both humans and ferrets. Like humans, ferrets showed spatially organized selectivity for different temporal and spectral modulation rates, that coarsely mimicked the types of selectivity we have previously observed in humans, replicating prior findings (Erb et al., 2019). And this selectivity was sufficiently similar that we could quantitatively predict response patterns to the synthetic sounds across species. These results do not imply that frequency and modulation tuning is the same across species, but do suggest that the organization is qualitatively similar.

Our results also do not imply that ferrets lack higher-order acoustic representations. Indeed, we found that ferrets’ spontaneous movements robustly discriminated between natural and synthetic ferret vocalizations, demonstrating behavioral sensitivity to the features which distinguish these sound sets, and this sensitivity was greater for ferret vocalizations than for either speech or music. But the manner in which species-relevant higher-order features are represented is likely distinct between humans and ferrets. Consistent with this idea, we found that differences between natural and synthetic sounds are weak, distributed throughout primary and non-primary regions, and show a mix of enhanced and suppressive responses (**Fig 4E**), unlike the strong, selective, and localized responses observed in human non-primary regions.

Our findings are broadly consistent with a recent study that compared responses to simple tone and noise stimuli between humans and macaques (Norman-Haignere et al., 2019). This study found that selective responses to tones vs. noise were larger in both primary and non-primary regions of human auditory cortex compared with macaques, which might reflect the importance of speech and music in humans where harmonic structure plays a central role. Our finding are unlikely to reflect greater tone selectivity because we have previously shown that non-primary regions respond preferentially to natural vs. temporally scrambled sounds with similar spectral properties (Norman-Haignere et al., 2015; Overath et al., 2015) (in addition we have found in pilot experiments that speech-selective regions respond strongly to whispered speech which lack tonal structure). Moreover, the prior study tested only two types of sounds (tones and noises) and thus was unable to broadly characterize how auditory representations differ between species. Here, we tested a wide and diverse range of natural and synthetic sounds that differ on many different ecologically relevant dimensions, and thus were able to compare the overall functional organization between humans and ferrets. As a consequence, we were able to identify a substantial divergence in neural representations at a specific point in the cortical hierarchy.

### Methodological advances

Our findings were enabled by a recently developed synthesis method, that makes it possible to synthesize sounds with frequency and modulation statistics that are closely matched to those in natural sounds (Norman-Haignere and McDermott, 2018). Because the synthetics are otherwise unconstrained, they lack higher-order acoustic properties present in complex natural sounds like speech and music (e.g. syllabic structure; musical notes, harmonies and rhythms). Comparing neural responses to natural and synthetic sounds thus provides a way to isolate responses to higher-order properties of natural stimuli that cannot be accounted for by modulation statistics. This methodological advance was critical to differentiating human and ferret cortical responses. Indeed, when considering natural or synthetic sounds alone, we observed very similar responses between species. We even observed selective responses to speech compared with other natural sounds in the ferret auditory cortex, due to the fact that speech has a unique range of spectrotemporal modulations. Thus, if we had only tested natural sounds, we might have concluded that speech and music-selective responses in the human non-primary auditory cortex reflect the same types of acoustic representations present in ferrets.

Our study illustrates the utility of wide-field imaging methods in comparing the brain organization of different species (Bimbard et al., 2018; Milham et al., 2018). Most animal physiology studies focus on measuring responses from single neurons or small clusters of neurons in a single brain region. While this approach is clearly essential to understanding the neural code at a fine grain, studying a single brain region can obscure larger-scale trends that are evident across the cortex. Indeed, if we had only measured responses in a single region of auditory cortex, we would have missed the most striking difference between humans and ferrets: the emergence of selective responses to natural sounds in non-primary regions of humans but not ferrets (**Fig 2E**).

Functional ultrasound imaging provides a powerful way of studying large-scale functional organization in small animals such as ferrets, since it has much better spatial resolution than fMRI (Macé et al., 2011; Bimbard et al., 2018). Because fUS responses are noisy, prior studies, including those from our own lab, have only been able to characterize responses to a single stimulus dimension, such as frequency, typically using a small stimulus set (Gesnik et al., 2017; Bimbard et al., 2018). Here, we developed a denoising method that made it possible to measure highly reliable responses to over a hundred stimuli in a single experiment. We were able to recover at least as many response dimensions as those detectable with fMRI and humans, and those response dimensions exhibited selectivity for a wide range of frequencies and modulation rates. Our study thus pushes the limits of what is possible using ultrasound imaging, and establishes fUS as an ideal method for studying the large-scale functional organization of the animal brain.

### Assumptions and limitations

The natural and synthetic sounds we tested were closely matched in their time-averaged cochlear frequency and modulation statistics, measured using a standard model of cochlear and cortical modulation tuning (Chi et al., 2005; Norman-Haignere and McDermott, 2018). We focused on time-averaged statistics because fMRI and fUS reflect time-averaged measures of neural activity, due to the temporally slow nature of hemodynamic responses. Thus, a similar response to natural and synthetic sounds indicates that the statistics being matched are sufficient to explain the voxel response. By contrast, a divergent voxel response indicates that the voxel responds to features of sound that are not captured by the model.

While divergent responses by themselves do not demonstrate a higher-order response, there are several reasons to think that the selectivity we observed in human non-primary regions is due to higher-order tuning. First, the fact that differences between natural and synthetic speech/music were much larger in non-primary regions clearly suggests that these differences are driven by higher-order processing above and beyond that present in primary auditory cortex, where spectrotemporal modulations appear to explain much of the voxel response. Second, the natural and synthetic sounds produced by our synthesis procedure are in practice closely matched on a wide variety on spectrotemporal filterbank models (Norman-Haignere and McDermott, 2018). As a consequence, highly divergent responses to natural and synthetic sounds rule out many such models. Third, the fact that responses were consistently larger for natural speech/music vs. synthetic speech/music suggests that these non-primary regions respond selectively to features in natural sounds that are not explicitly captured by spectrotemporal modulations and are thus absent from the synthetic sounds.

As with any study, our conclusions are limited by the precision and coverage of our neural measurements. For example, fine-grained temporal codes, which have been suggested to play an important role in vocalization encoding (Schnupp et al., 2006), cannot be detected with fUS. However, we note that the resolution of fUS is substantially better than fMRI, particularly in the spatial dimension (voxel sizes were more than 1000 times smaller) and thus the species differences we observed are unlikely to be explained by differences in the resolution of fUS vs. fMRI. It is also possible that ferrets might show more prominent differences between natural and synthetic sounds outside of auditory cortex. But even if this were true, it would still demonstrate a clear species difference because humans show robust selectivity for natural sounds in non-primary regions just outside of primary auditory cortex, while ferrets evidently do not.

### Possible nature and causes of differences in higher-order selectivity

What features might non-primary human auditory cortex represent, given that spectrotemporal modulations do not explain all of the response? Although these regions respond selectively to speech and music, they are not driven by semantic meaning or explicit musical training (Overath et al., 2015; Boebinger et al., 2020), are located just beyond primary auditory cortex, and show evidence of having short integration periods on the scale of hundreds of milliseconds (Overath et al., 2015). This pattern suggests nonlinear selectivity for short-term temporal and spectral structure present in speech syllables or musical notes (e.g. harmonic structure, pitch contours, and local periodicity). This hypothesis is consistent with recent work showing sensitivity to phonotactics in non-primary regions of the superior temporal gyrus (Leonard et al., 2015; Brodbeck et al., 2018; Di Liberto et al., 2019), and with a recent study showing that deep neural networks trained to perform challenging speech and music tasks are better able to predict responses in non-primary regions of human auditory cortex (Kell et al., 2018).

Why don’t we observe similar neural selectivity in ferrets for vocalizations? Ferret vocalizations clearly exhibit additional structure not captured by spectrotemporal modulations, since the animals showed large and spontaneous increases in motion for natural vs. synthetic vocalizations. This increase in motion was greater for vocalizations than for either speech or music, clearly reflecting the behavioral significance of vocalizations to ferrets. However, this additional structure may play a less-essential role in their everyday hearing compared with that of speech and music in humans. Other animals that depend more on higher-order acoustic representations might show more human-like selectivity in non-primary regions. For example, marmosets have a relatively complex vocal repertoire (Agamaite et al., 2015) and depend more heavily on vocalizations than many other species (Eliades and Miller, 2017), and thus might exhibit more prominent selectivity for higher-order properties in their calls. It may also be possible to experimentally enhance selectivity for higher-order properties via extensive exposure and training, particularly at an early age of development (Polley et al., 2006; Srihasam et al., 2014). All of these questions could be addressed in future work using the methods developed here.

## Methods

### Animal preparation

Experiments were performed in two head-fixed awake ferrets (A and T), across one or both hemispheres (Study 1: A_left_, A_right_, T_left_, T_right_; Study 2: A_left_, T_left_, and T_right_). Ferret A was a mother (had one litter of pups), while ferret T was a virgin. Experiments were approved by the French Ministry of Agriculture (protocol authorization: 21022) and strictly comply with the European directives on the protection of animals used for scientific purposes (2010/63/EU). Animal preparation and fUS imaging were performed as in Bimbard et al. (2018). Briefly, a metal headpost was surgically implanted on the skull under anaesthesia. After recovery from surgery, a craniotomy was performed over auditory cortex and then sealed with an ultrasound-transparent Polymethylpentene (TPX™) cover, embedded in an implant of dental cement. Animals could then recover for one week, with unrestricted access to food, water and environmental enrichment. Imaging windows were maintained across weeks with appropriate interventions when tissue and bone regrowth were shadowing brain areas of interest.

### Ultrasound imaging

fUS data are collected as a series of 2D images or ‘slices’. Slices were collected in the coronal plane and were spaced 0.4 mm apart. The slice plane was varied across sessions in order to cover the region-of-interest which included both primary and non-primary regions of auditory cortex. One or two sessions were performed on each day of recording. The resolution of each voxel was 0.1 x 0.1 x ∼0.4 mm (the latter dimension, called elevation, being slightly dependent on the depth of the voxel). The overall voxel volume (0.004 mm^3^) was more than a thousand times smaller than the voxel volume used in our human study (which was either 8 or 17.64 mm^3^ depending on the subjects/paradigm), which helps to account for their smaller brain.

A separate “Power Doppler” image/slice was acquired every second. Each of these images was computed by first collecting 300 sub-images or ‘frames’ in a short 600 ms time interval (500 Hz sampling rate). Those 300 frames were then filtered to discard global tissue motion from the signal (Demené et al., 2015) (the first 55 principal components were discarded because they mainly reflect motion; see Demené et al., 2015 for details). The blood signal energy also known as Power Doppler was computed for each voxel by summing the squared magnitudes across the 300 frames separately for each pixel (Macé et al., 2011). Power Doppler is approximately proportional to blood volume (Macé et al., 2011).

Each of the 300 frames was itself computed from 11 tilted plane wave emissions (−10° to 10° with 2° steps) fired at a pulse repetition frequency of 5500 Hz. Frames were reconstructed from these plane wave emissions using an in-house, GPU-parallelized delay-and-sum beamforming algorithm (Macé et al., 2011).

### Stimuli for Experiment I

We tested 40 natural sounds: 36 sounds from our prior experiment plus 4 ferret vocalizations (fight call, pup call, fear vocalization, and play call). Each natural sound was 10 seconds in duration. For each natural sound, we synthesized four synthetic sounds, matched on a different set of acoustic statistics of increasing complexity: cochlear, temporal modulation, spectral modulation, and spectrotemporal modulation. The modulation-matched synthetics were also matched in their cochlear statistics to ensure that differences between cochlear and modulation-matched sounds must be due to the addition of modulation statistics. The natural and synthetic sounds were identical to those in our prior paper, except for the four additional ferret vocalizations, which were synthesized using the same algorithm. We briefly review the algorithm below.

Cochlear statistics were measured from a cochleagram representation of sound, computed by convolving the sound waveform with filters designed to mimic the pseudo-logarithmic frequency resolution of cochlear responses (McDermott and Simoncelli, 2011). The cochleagram for each sound was composed of the compressed envelopes of these filter responses (compression is designed to mimic the effects of cochlear amplification at low sound levels). Modulation statistics were measured from filtered cochleagrams, computed by convolving each cochleagram in time and frequency with a filter designed to highlight modulations at a particular temporal rate and/or spectral scale (Chi et al., 2005). The temporal and spectral modulation filters were only modulated in time or frequency, respectively. There were 9 temporal filters (best rates: 0.5, 1, 2, 4, 8, 16, 32, 64, and 128 Hz) and 6 spectral filters (best scales: 0.25, 0.5, 1, 2, 4, 8 cycles per octave). Spectrotemporal filters were created by taking the outer-product of all pairs of temporal and spectral filters in the 2D fourier domain, which results in oriented gabor-like filters.

Our synthesis algorithm matches time-averaged statistics of the cochleagrams and filtered cochleagrams via a histogram-matching procedure that implicitly matches all time-averaged statistics of the responses (separately for each frequency channel of the cochleagrams and filtered cochleagrams). This choice is motivated by the fact that both fMRI and fUS reflect time-averaged measures of neural activity, because the temporal resolution of hemodynamic changes is much slower than the underlying neuronal activity. As a consequence, if the fMRI or fUS response is driven by a particular set of acoustic features, we would expect two sounds with similar time-averaged statistics for those features to yield a similar response. We can therefore think of the natural and synthetic sounds as being matched under a particular model of the fMRI or fUS response (a more formal derivation of this idea is given in Norman-Haignere et al., 2018).

We note that the filters used to compute the cochleagram were designed to match the frequency resolution of the human cochlea, which is thought to be somewhat finer than the frequency resolution of the ferret cochlea (Walker et al., 2019). In general, synthesizing sounds from broader filters results in synthetics that differ slightly more from the originals. And thus if we had used cochlear filters designed to mimic the frequency tuning of the ferret cochlea, we would expect the cochlear-matched synthetic sounds to differ slightly more from the natural sounds. However, given that we already observed highly divergent responses to natural and cochlear-matched synthetic sounds in both species, it is unlikely that using broader cochlear filters would change our findings. In general, we have found the matching procedure is not highly sensitive to the details of the filters used. For example, we have found that sounds matched on the spectrotemporal filters used here and taken from Chi et al. (2005), are also well matched on filters with half the bandwidth, with phases that have been randomized, and with completely random filters (Norman-Haignere and McDermott, 2018).

### Stimuli for Experiment II

Experiment II tested a larger set of 30 ferret vocalizations (5 fight calls, 17 single-pup calls, and 8 multi-pup calls where the calls from different pups overlapped in time). The vocalizations consisted of recordings from several labs (our own, Stephen David’s and Andrew King’s laboratories). For comparison, we also tested 14 speech sounds and 16 music sounds, yielding 60 natural sounds in total. For each natural sound, we created a synthetic sound matched on the full spectrotemporal model. We did not synthesize sounds for the sub-models (cochlear, temporal modulation, and spectral modulation), since our goal was to test if there were divergent responses to natural and synthetic ferret vocalizations for spectrotemporally-matched sounds, like those present in human non-primary auditory cortex for speech and music sounds.

### Procedure for presenting stimuli

Sounds were played through calibrated earphones (Sennheiser IE800 earphones, HDVA 600 amplifier, 65 dB) while recording hemodynamic responses via fUS imaging. In our prior fMRI experiments in humans, we had to chop the 10 second stimuli into 2-second excerpts in order to present the sounds in between scan acquisitions, because MRI acquisitions produce a loud sound that would otherwise interfere with hearing the stimuli. Because fUS imaging produces no audible noise, we were able to present the entire 10 second sound without interruption. The experiment was composed of a series of 20-second trials, and fUS acquisitions were synchronized to trial onset. On each trial, a single 10-second sound was played, with 7 seconds of silence before the sound to establish a response baseline, and 3 seconds of post-stimulus silence to allow the response to return to baseline. There was a randomly chosen 3 to 5 second gap between each trial. Sounds were presented in random order, and each sound was repeated 4 times.

### Mapping of tonotopic organization with pure tones

Tonotopic organization was assessed using previously described methods (Bimbard et al., 2018). In short, responses were measured to 2-second long pure tones from 5 different frequencies (602 Hz, 1430 Hz, 3400 Hz, 8087 Hz, 19234 Hz). The tones were played in random order, with 20 trials/frequency. Data was denoised using the same method described in *Denoising Part I: Removing components outside of cortex*. Tonotopic maps were created by determining the best frequency of each voxel, defined as the tone evoking the largest Power Doppler signal. We then used these functional landmarks in combination with brain and vascular anatomy to establish the borders between primary and non-primary areas in all hemispheres, as well as to compare them to those obtained with natural sounds (see **Fig S4A**).

### Brain map display

Views from above were obtained by computing the average of the variable of interest in each vertical column of voxels from the upper part of the manually defined cortical mask. This is justified by the fact that measures were coherent across depth (see **Fig S4** for examples). However, we note that having a three-dimensional view prevents us from missing specific responsive areas sometimes buried in the depth of the sulci.

### Normalized Squared Error (NSE) maps

Like fMRI, the response timecourse of each fUS voxel shows a gradual build-up of activity after a stimulus, due to the slow and gradual nature of blood flow changes. The shape of this response timecourse is similar across different sounds, but the magnitude varies (**Fig 1C**) (fMRI responses show the same pattern). We therefore measured the response magnitude of each voxel by averaging the response to each sound across time (from 3 to 11 seconds post-stimulus onset), yielding one number per sound. Responses were measured from denoised data. We describe the denoising procedure at the end of the Methods because it is more involved than our other analyses.

We compared the response magnitude to natural and corresponding synthetic sounds using the normalized squared error (NSE), the same metric used in humans. The NSE takes a value of 0 if the response to natural and synthetic sounds is identical, and 1 if there is no correspondence between responses to natural and synthetic sounds. The NSE is defined as:

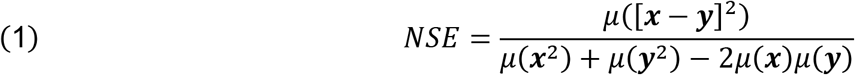

where ***x*** and ***y*** are response vectors across the sounds being compared (i.e. natural and synthetic) and *μ*(.) indicates the vector mean. We noise-corrected the NSE using the test-retest reliability of the voxel responses (see Norman-Haignere et al., 2018 for details). However, we measured the NSE from denoised data, which was highly reliable, and our correction procedure thus only had a small effect on the resulting values.

### Annular ROI analyses

We used the same annular ROI analyses from our prior paper to quantify the change in NSE values (or lack thereof) across the cortex. We binned voxels based on their distance to the center of primary auditory cortex, defined tonotopically. We used smaller bin sizes in ferrets (0.5 mm) than humans (5 mm) due to their smaller brains (results were not sensitive to the choice of bin size). **Figure 2F** plots the median NSE value in each bin, plotted separately for each human subject and for each hemisphere of each ferret. To statistically compare different models (e.g. cochlear vs. spectrotemporal), we averaged the NSE values across all bins and hemispheres/subjects separately for each model, bootstrapped the resulting statistics by resampling across the sound set (1000 times), and counted the fraction of samples that overlapped between models (multiplying by 2 to arrive at a two-sided p-value). To compare species, we measured the slope of the NSE vs. distance curve separately for each hemisphere/animal. We found that the slope in every hemisphere of every ferret was less than the slope of every hemisphere of every human subject, which is significant with a sign test (p < 0.01; for each ferret hemisphere there were 8 human subjects to compare with).

### Component analyses

To investigate the organization of fUS responses to the sound set, we applied the same voxel decomposition used in our prior work in humans to identify a small number of component response patterns that explained a large fraction of the response variation. Like all factorization methods, each voxel is modeled as the weighted sum of a set of canonical response patterns that are shared across voxels. The decomposition algorithm is similar to standard algorithms for independent component analysis (ICA) in that it identifies components that have a non-Gaussian distribution of weights across voxels by minimizing the entropy of the weights (the Gaussian distribution has the highest entropy of any distribution with fixed variance). This optimization criterion is motivated by the fact that independent variables become more Gaussian when they are linearly mixed, and non-Gaussianity thus provides a statistical signature that can be used to unmix the latent variables. Our algorithm differs from standard algorithms for ICA in that it estimates entropy using a histogram, which is effective if there are many voxels, as is the case with fMRI and fUS (40882 fUS voxels for experiment I, 38366 fUS voxels for experiment II).

We applied our analyses to the denoised response timecourse of each voxel across all sounds (each column of the data matrix contained the concatenated response timecourse of one voxel across all sounds). Our main analysis was performed on voxels concatenated across both animals tested. The results however were similar when the analysis was performed on data from each animal. The number of components was determined via a cross-validation procedure described in the section on denoising.

We examined the inferred components by plotting and comparing their response profiles to the natural and synthetic sounds, as well as plotting their anatomical weights in the brain. We also correlated the response profiles across all sounds with measures of cochlear and spectrotemporal modulation energy. Cochlear energy was computed by averaging the cochleagram for each sound across time. Spectrotemporal modulation energy was calculated by measuring the strength of modulations in the filtered cochleagrams (which highlight modulations at a particular temporal rate and/or spectral scale). Modulation strength was computed as the standard deviation across time of each frequency channel of the filtered cochleagram. The channel-specific energies were then averaged across frequency, yielding one number per sound and spectrotemporal modulation rate.

We used a permutation test across the sound set to assess the significance of correlations with frequency and modulation features. Specifically, we measured the maximum correlation across all frequencies and all modulation rates tested, and we compared these values with those from a null distribution computed by permuting the correspondence across sounds between the features and the component responses (1000 permutations). We counted the fraction of samples that overlapped the null distribution and multiplied by two in order to arrive at a two-sided p-value. For every component, we found that correlations with frequency and modulation features were significant (p < 0.01).

### Predicting human components from ferret responses

To quantify which component response patterns were shared across species, we tried to linearly predict components across species (**Fig S6/S7**). Each component was defined by its average response to the 36 natural and corresponding synthetic sounds, matched on the full spectrotemporal model. We attempted to predict each human component from all of the ferret components and vice versa, using cross-validated ridge regression (9 folds). The ridge parameter was chosen using nested cross-validation within the training set (also 9 folds; testing a wide range from 2^-100^ to 2^100^). Each fold contained pairs of corresponding natural and synthetic sound, so that there would be no overlap between the train and test sounds.

For each component, we separately measured how well we could predict the response to synthetic sounds (**Fig S6B/S7A**) – which isolates selectivity for frequency and modulation statistics present in natural sounds – as well as how well we could predict the difference between responses to natural vs. synthetic sounds (**Fig S6C/FigS7B**) – which isolates selectivity for features in natural sounds that are not explained by frequency and modulation statistics. We quantified prediction accuracy using the noise-corrected NSE, and we used (1 − *NSE*). ^2 as a measure of explained variance. This choice is motivated by the fact (1 − *NSE*) is equivalent to the Pearson correlation for signals with equal mean and variance and thus (1 − *NSE*). ^2 is analogous to the squared Pearson correlation, which is a standard measure of explained variance.

We multiplied these explained variance estimates by the total response variance of each component for either synthetic sounds or for the difference between natural and synthetic sounds (**Fig S6D/Fig S7C** shows the total variance alongside the fraction of that total variance explained by the cross-species prediction). We noise-corrected the total variance using the equation below:

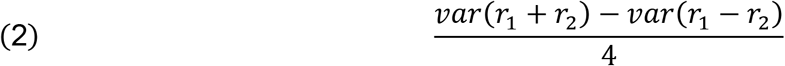

where *r*_1_ and *r*_2_ are two independent response measurements. Below we give a brief derivation of this equation, where *r*_1_ and *r*_2_ are expressed as the sum of a shared signal (*s*) that is repeated across measurements plus independent noise (*n*_1_ and *n*_2_) which is not. This derivation utilizes the fact that the variance of independent signals that are summed or subtracted is equal to the sum of their respective variances.

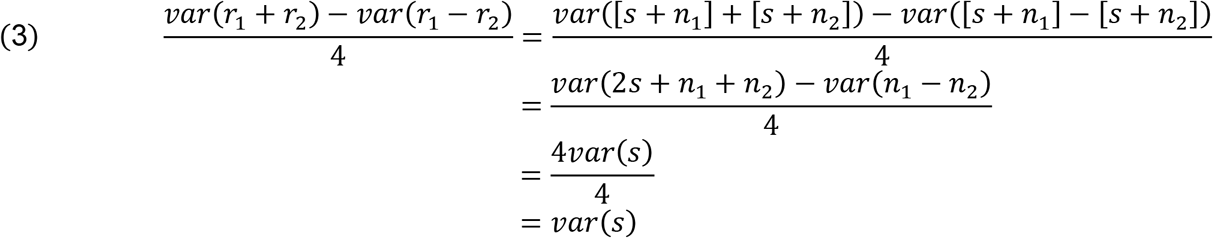

The two independent measurements used for noise correction were derived from different human or ferret subjects. The measurements were computed by attempting to predict group components from each individual subject using the same cross-validated regression procedure described above. The two measurements in ferrets came from the two animals tested (A and T). And the two measurements in humans came from averaging across two non-overlapping sets of subjects (4 in each group; groups chosen to have similar SNR).

For this analysis, the components were normalized so that the RMS magnitude of their weights was equal. As a consequence, components that explained more response variance also had larger response magnitudes. We also adjusted the total variance across all components to equal 1.

#### Comparing the similarity of natural and synthetic sounds from different categories

We computed maps showing the average difference between natural and synthetic sounds from different categories (**Fig 4E**). So that the scale of the differences could be compared across species, we divided the measured differences by the standard deviation of each voxel’s response across all sounds. We also separately measured the NSE for sounds from different categories (**Fig 4F**,**G**). The normalization term in the NSE equation (denominator of equation 1) was averaged across all sounds in order to ensure that the normalization was the same for all sounds/categories and thus that we were not inadvertently normalizing-away meaningful differences between the sounds/categories.

### Denoising Part I: Removing components outside of cortex

Ultrasound responses in awake animals are noisy, which has limited its usage to mapping simple stimulus dimensions (e.g. frequency) where a single stimulus can be repeated many times (Bimbard et al., 2018). To overcome this issue, we developed a denoising procedure that substantially increased the reliability of the voxel responses (**Fig S9**). The procedure had two parts. The first part, which is described in this section, removed prominent signals outside of cortex, which are likely to reflect movement or other sources of noise. The second part enhanced reliable signals. Code implementing the denoising procedures will be made available upon publication.

We separated voxels into those inside and outside of cortex, since responses outside of the cortex by definition do not contain stimulus-driven cortical responses, but do contain sources of noise like motion. We then used canonical correlation analysis (CCA) to find a set of response timecourses that were robustly present both inside and outside of cortex, since such timecourses are both likely to reflect noise and likely to distort the responses-of-interest. We projected-out the top 20 canonical components (CCs) from the data set, which we found scrubbed the data of motion-related signals (**Fig S9A**; motion described below).

This analysis was complicated by one key fact: the animals reliably moved more during the presentation of some sounds (**Fig 4C**). Thus, noise-induced activity outside-of-cortex is likely to be correlated with sound-driven neural responses inside-of-cortex, and removing CCs will thus remove both noise and genuine sound-driven activity. To overcome this issue, we took advantage of the fact that sound-driven responses will by definition be reliable across repeated presentations of the same sound, while motion-induced activity will vary from trial-to-trial for the same sound. We thus found canonical components where the residual activity after removing trial-averaged responses was shared between responses inside and outside of cortex, and we then removed the contribution of these components from the data. We give a detailed description and motivation of this procedure in the **Appendix**, and show the results of a simple simulation demonstrating its efficacy.

To assess the effect of this procedure on our fUS data, we measured how well it removed signals that were correlated with motion (**Fig S9A**). Motion was measured using a video recording of the animals’ face. We measured the motion energy in the video as the average absolute deviation across adjacent frames, summed across all pixels. We correlated this motion timecourse with the residual timecourse of every voxel after subtracting off trial-averaged activity. **Figure S9A** plots the mean absolute correlation value across voxels as a function of the number of canonical components removed (motion can induce both increased and decreased fUS signal and thus it was necessary to take the absolute value of the correlation before averaging). We found that removing the top 20 CCs substantially reduced motion correlations.

We also found that removing the top 20 CCs removed the spatial striping in the voxel responses, which is a stereotyped feature of motion due to the interaction between motion and blood vessels. To illustrate this effect, **Figure S9B** shows the average difference between responses to natural vs. synthetic sounds in Experiment II (vocalization experiment). Before denoising, this difference map shows a clear striping pattern likely due to the fact that the animals moved more during the presentation of the natural vs. synthetic sounds. The denoising procedure largely eliminated this striping pattern.

### Denoising Part II: Enhancing signal using DSS

After removing components likely to be driven by noise, we applied a second procedure designed to enhance reliable components in the data. Our procedure is a variant of a method that is often referred to as “denoising source separation” (DSS) or “joint decorrelation” (de Cheveigné and Parra, 2014). In contrast with principal component analysis (PCA), which finds components that have high variance, DSS emphasizes components that have high variance after applying a “biasing” operation that is designed to enhance some aspect of the data. The procedure begins by whitening the data such that all response dimensions have equal variance, the biasing operation is applied, and PCA is then used to extract the components with highest variance after biasing. In our case, we biased the data to enhance response components that were reliable across stimulus repetitions and across the slices from all animals. We note that unlike fMRI, data from different slices come from different sessions. As a consequence, the noise from different slices will be independent. Thus, any response components that are consistent across slices and animals are likely to reflect true, stimulus-driven responses.

The input to our analysis was a set of matrices. Each matrix contained data from a single stimulus repetition and slice. Only voxels from inside of cortex were analyzed. Each column of each matrix contained the response timecourse of one voxel to all of the sounds (concatenated), denoised using the procedure described in Part I. The response of each voxel was converted to units of percent signal change (the same units used for fMRI analyses) by subtracting and dividing by the pre-stimulus period (also known as percent Cerebral Blood Volume or %CBV in the fUS literature).

Our analysis involved five steps:

1. We whitened each matrix individually.
2. We averaged the whitened response timecourses across repetitions, thus enhancing responses that are reliable across repetitions.
3. We concatenated the repetition-averaged matrices for all slices across the voxel dimension, thus boosting signal that is shared across slices and animals.
4. We extracted the top N principal components (PCs) with the highest variance from the concatenated data matrix. The number of components was selected using cross-validation (described below). Because the matrices for each individual repetition and slice have been whitened, the PCs extracted in this step will *not* reflect the components with highest variance, but will instead reflect the components that are the most reliable across repetitions and across slices/animals. We thus refer to these components as “reliable components” (*R*).
5. We then projected the data onto the top N reliable components (*R*):

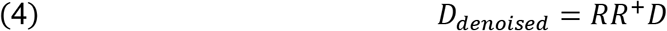

where *D* is the denoised response matrix from Part I.

We used cross-validation to test the efficacy of this denoising procedure and select the number of components (**Fig S2**).

The analysis involved the following steps:

1. We divided the sound set into training (75%) and test (25%) sounds. Each set contained corresponding natural and synthetic sounds so that there would be no overlap between train and test sets. We attempted to balance the train and test sets across categories, such that each split had the same number of sounds from each category.
2. Using responses to just the train sounds (*D*_*train*_), we computed reliable components (*R*_*train*_) using the procedure just described (steps 1-4).
3. We calculated voxel weights for these components:

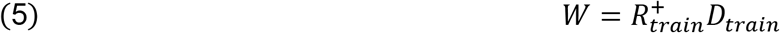
4. We used this weight matrix, which was derived entirely from train data, to denoise responses to the test sounds:

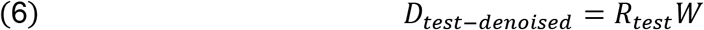

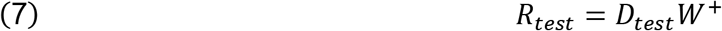

To evaluate whether the denoising procedure improved predictions, we measured responses to the test sound set using two independent splits of data (odd or even repetitions). We then correlated the responses across the two splits either before or after denoising.

**Figure S2A** plots the split-half correlation of each voxel before vs. after denoising for every voxel in cortex (using an 8-component model). For this analysis, we either denoised one split of data (blue dots) or both splits of data (green dots). Denoising one split provides a fairer test of whether the denoising procedure enhances SNR, while denoising both splits demonstrates the overall boost in reliability. We also plot the upper bound on the split-half correlation when denoising one split of data (black line), which is given by the square root of the split-half reliability of the original data. We found that our denoising procedure substantially increased reliability with the denoised-correlations remaining close to the upper bound. When denoising both splits, the split-half correlations were close to 1, indicating a highly reliable response.

**Figure S2B** plots a map in one animal of the split-half correlations when denoising one split of data along with a map of the upper bound. As is evident, the denoised correlations remain close to the upper bound throughout primary and non-primary auditory cortex.

**Figure S2C** shows the median split-half correlation across voxels as a function of the number of components. Performance was best using ∼8 components in both experiments.

## Acknowledgements

We thank Sophie Bagur for careful reading of the manuscript and precious comments. This work was supported by ANR-17-EURE-0017 and the NVIDIA Corporation (NVidia GPU Grant program), an AMX doctoral fellowship to AL, CDSN doctoral fellowship and EMBO ALTF 740-2019 postdoctoral fellowship to CB, grants from National Institutes of Health (NIDCD, DC005779) and ERC Advanced grant 787836-NEUME to SAS, postdoctoral grants to SNH from the HHMI (through the Life Sciences Research Foundation) and the NIH (K99/R00 award), ANR-10-IDEX-0001-02 and ANR-JCJC-DynaMiC to YB.

## Author contributions

AL, CB and YB designed the experiments. SNH designed sound stimuli. CD provided technical support. AL and CB recorded the data. AL, CB, SNH and YB designed and carried out the analyses and statistical testing. CD and SAS provided feedback on the manuscript. AL, CB, SNH and YB wrote all aspects of the manuscript.

---

**Table S1:**
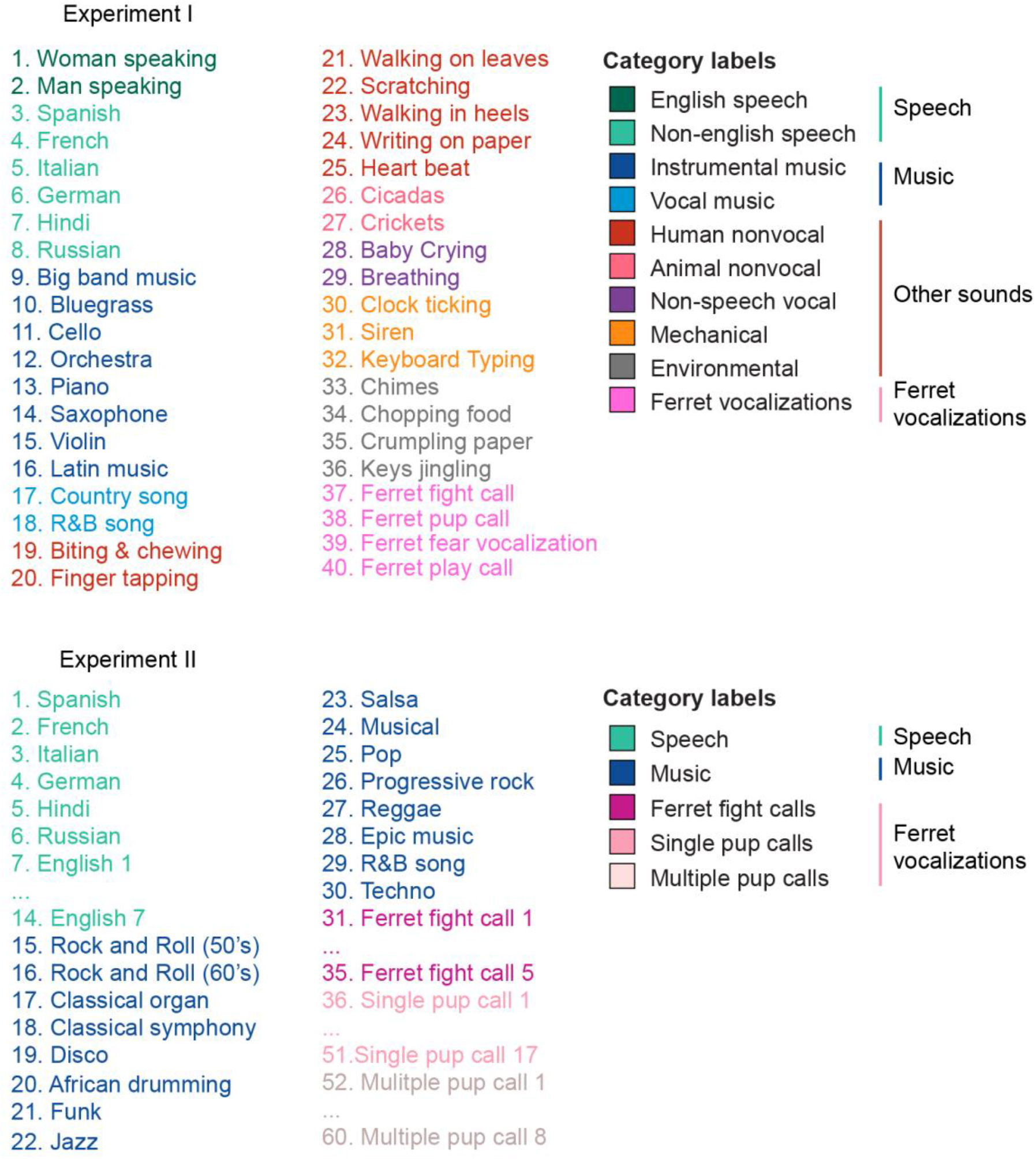
List of sounds used in both experiments. Names of sounds used in Experiments I and II, grouped by category at both fine and coarse scales.

## Supplementary information

**Figure S1.**
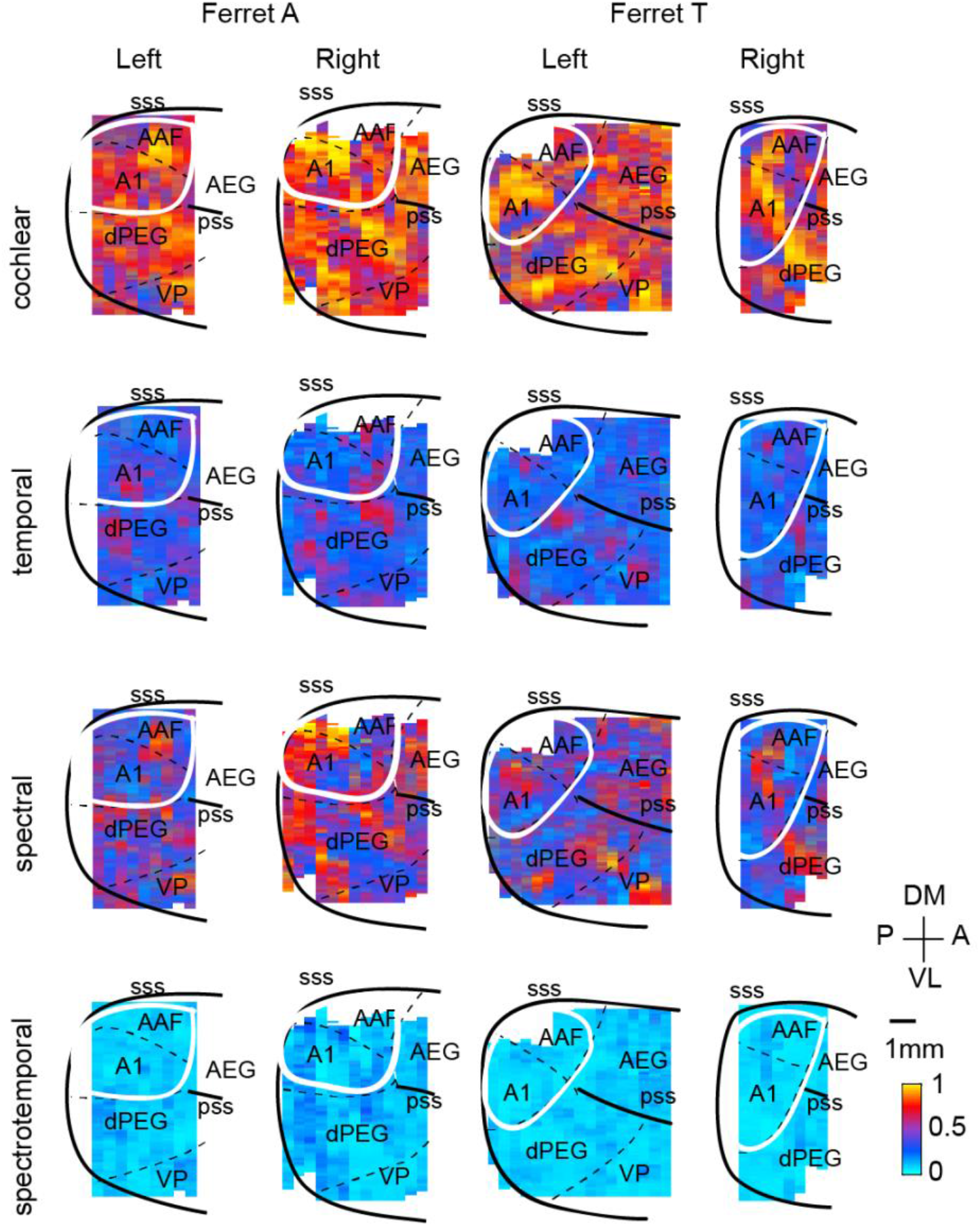
Dissimilarity maps for all hemispheres and animals. Same format as Figure 2E.

**Figure S2.**
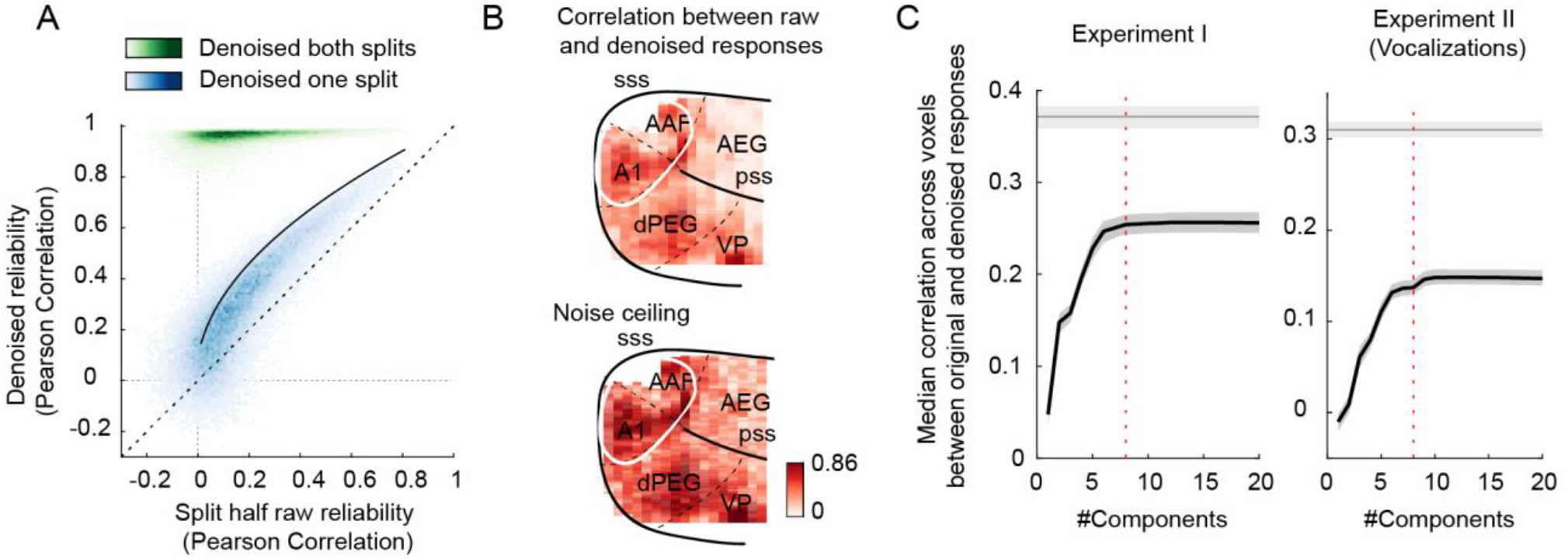
The effect of enhancing reliable signal using a procedure similar to “DSS”. (see Denoising Part II in Methods) (de Cheveigné and Parra, 2014). **A**, Voxel responses were denoised by projecting their timecourse onto components that were reliably present across repetitions, slices and animals. This figure plots the test-retest correlation across independent splits of data before (x-axis) and after (y-axis) denoising (data from Experiment I). Each dot corresponds to a single voxel. We denoised either one split of data (blue dots) or both splits of data (green dots). Denoising one split provides a fairer test of whether the denoising procedure enhances SNR. Denoising both splits shows the overall effect on response reliability. The theoretical upper-bound for denoising one split of data is shown by the black line. The denoising procedure substantially increased data reliability, with the one-split correlations hugging the upper-bound. This plot shows results from an 8-component model. **B**, This figure plots split-half correlations for denoised data (one split) as a map (upper panel), along with a map showing the upper bound (right). Denoised correlations were close to their upper bound throughout auditory cortex. **C**, This figure plots the median denoised correlation across voxels (one split) as a function of the number of components used in the denoising procedure. Gray line plots the upper bound. Shaded areas correspond to 95% confidence interval, computed via bootstrapping across the sound set. Results are shown for both Experiments I (left) and II (right). Predictions were near their maximum using ∼8 components in both experiments (the 8-component mark is shown by the vertical dashed line).

**Figure S3.**
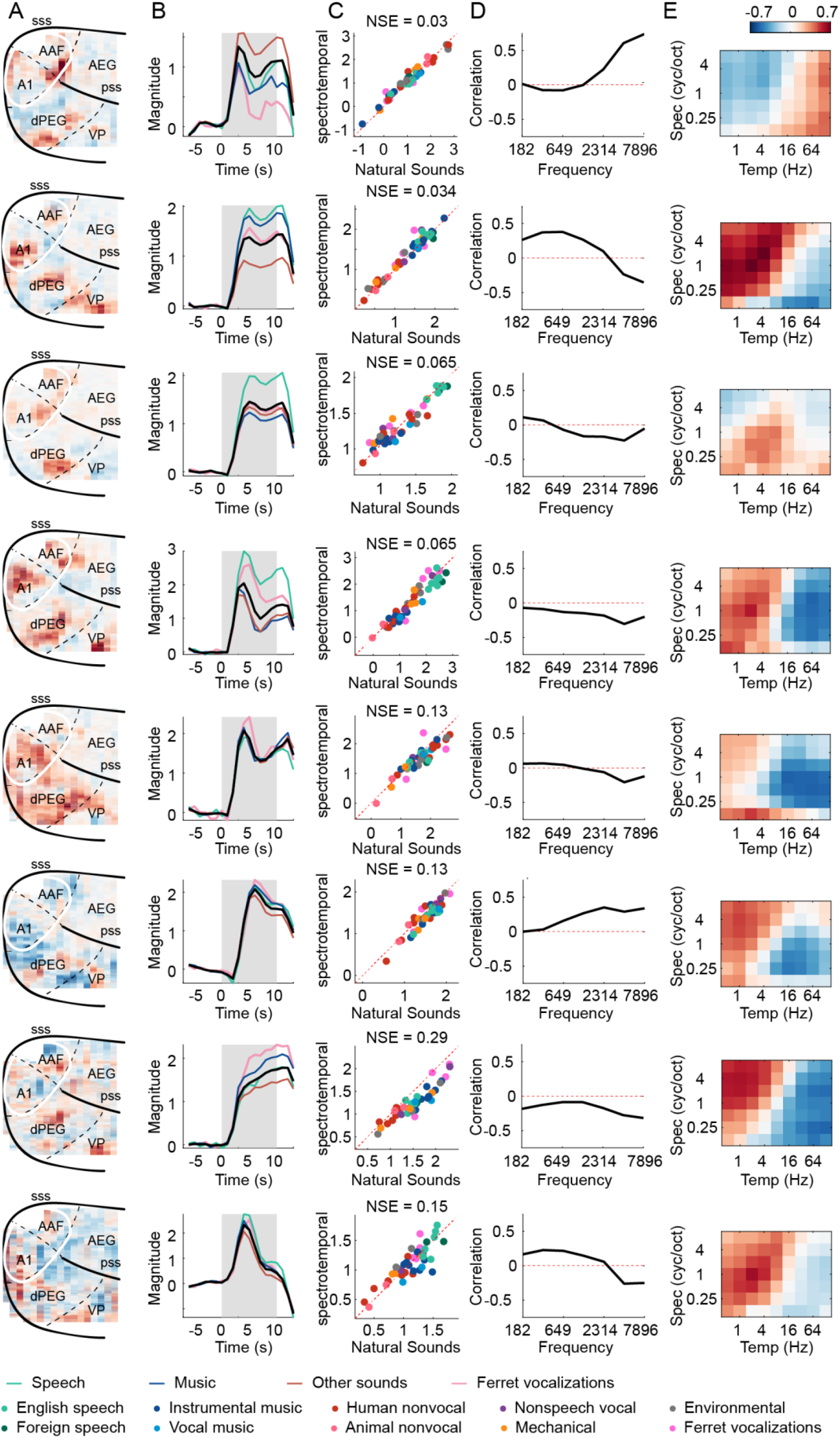
Results from all 8 ferret components. Same format as **Figure 3**, except for panel **B**, which plots the temporal response of the components. Black line shows the average across all natural sounds. Colored lines correspond to major categories (see **Table S1**): speech (green), music (blue), vocalizations (pink) and other sounds (brown). Note that the temporal shape varies across components, but is very similar across sounds/categories within a component, which is why we summarized component responses by their time-averaged response to each sound.

**Figure S4.**
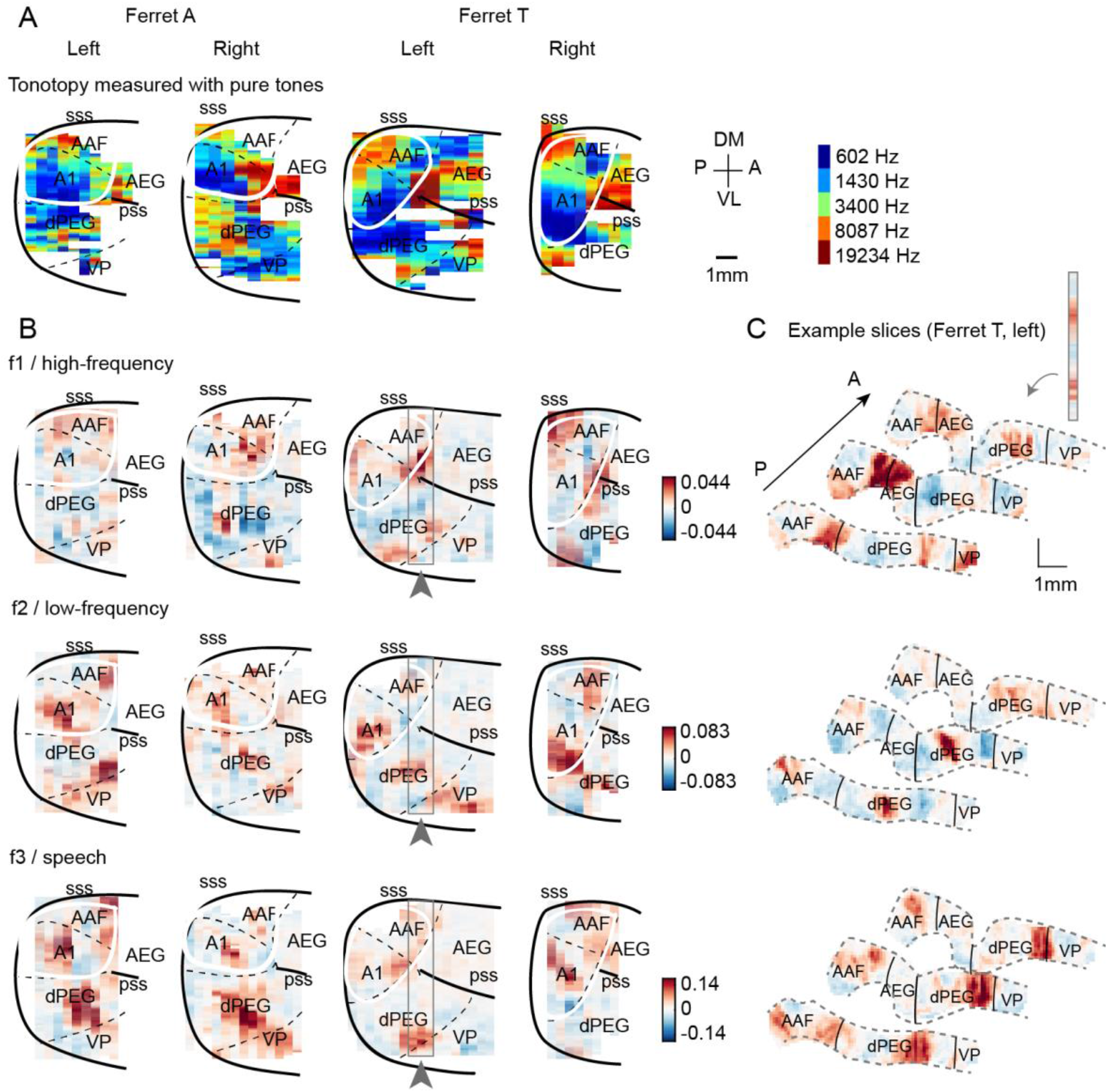
Component weight maps from all hemispheres and ferrets. **A**, For reference with the weight maps in panel B, tonotopic maps measured using pure tones are shown for all hemispheres. **B**, Voxel weight maps from the three components shown in **Figure 3** for all hemispheres of all ferrets tested. **C**, Voxel weights for three example coronal slices from Ferret T, left hemisphere. Grey outlines in panel B indicate their location in the “surface” view. Each slice corresponds to one vertical strip from the maps in panel B. The same slices are shown for all three components.

**Figure S5.**
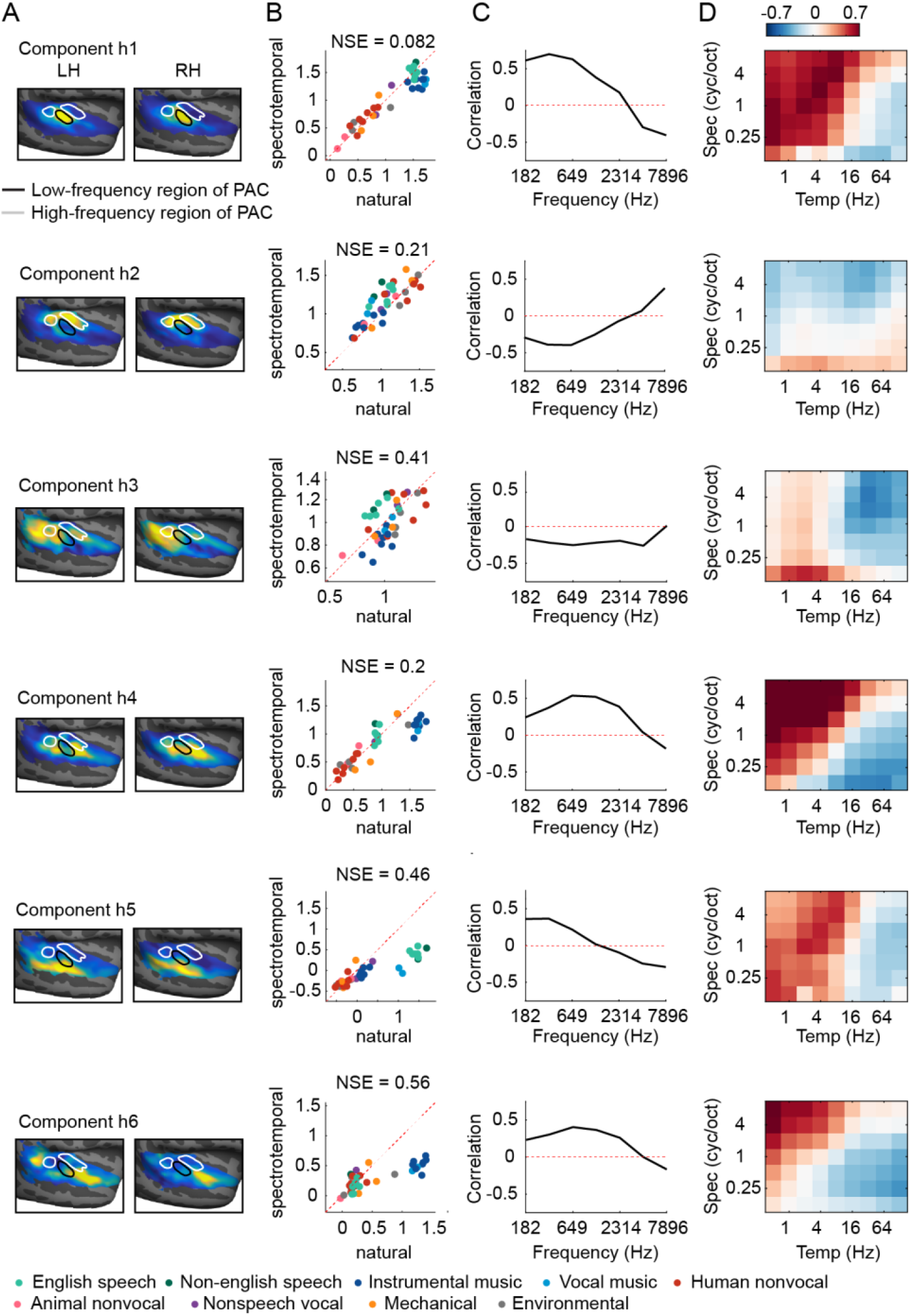
Human components. This figure shows the anatomy and response properties of the six human components inferred in prior work (Norman-Haignere et al., 2015; Norman-Haignere and McDermott, 2018). Same format as **Figure 3**, which plots ferret components. Weight maps (panel A) plot group-averaged maps across subjects.

**Figure S6.**
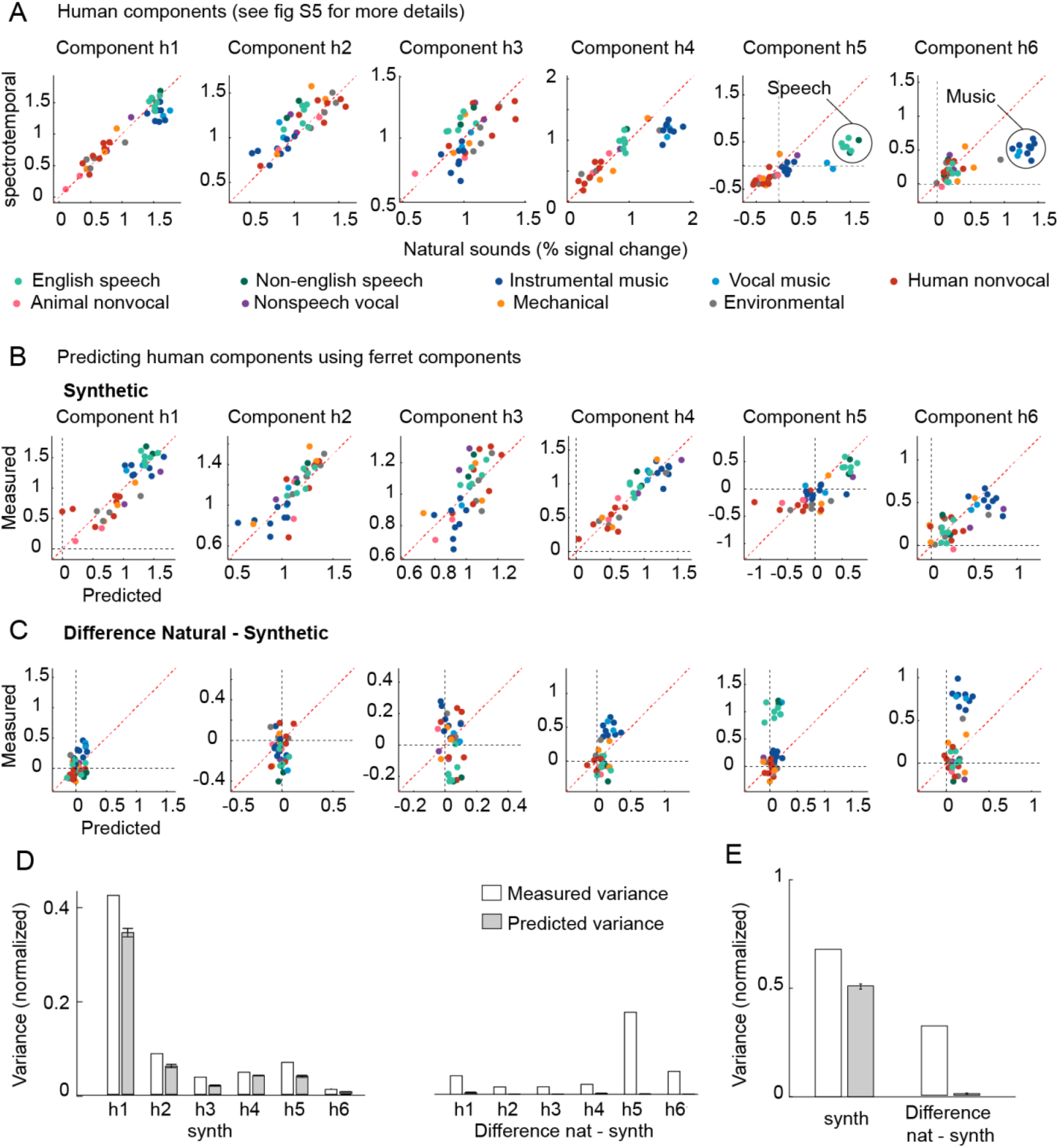
Predicting human component responses from ferrets. This figure plots the results of trying to predict the six human components inferred from our prior work (Norman-Haignere et al., 2015; Norman-Haignere and McDermott, 2018) from the eight ferret components inferred here (see **Fig S7** for the reverse). **A**, For reference, the response of the six human components to natural and spectrotemporally matched synthetic sounds is re-plotted here. Components h1-h4 produced similar responses to natural and synthetic sounds, and had weights that clustered in and around primary auditory cortex (**Fig S5**). Components h5 and h6 responded selectively to natural speech and natural music, respectively, and had weights that clustered in non-primary regions. **B**, This panel plots the measured response of each human component to spectrotemporally matched synthetic sounds, along with the predicted response from ferrets. **C**, This panel plots the difference between responses to natural and spectrotemporally-matched synthetic sounds along with the predicted difference from the ferret components. **D**, Plots the total response variance (white bars) of each human component to synthetic sounds (left) and to the difference between natural and synthetic sounds (right) along with the fraction of that total response variance predictable from ferrets (gray bars) (all variance measures are noise-corrected). Error bars show the 95% confidence interval, computed via bootstrapping across the sound set. **E**, Same as D, but averaged across components.

**Figure S7.**
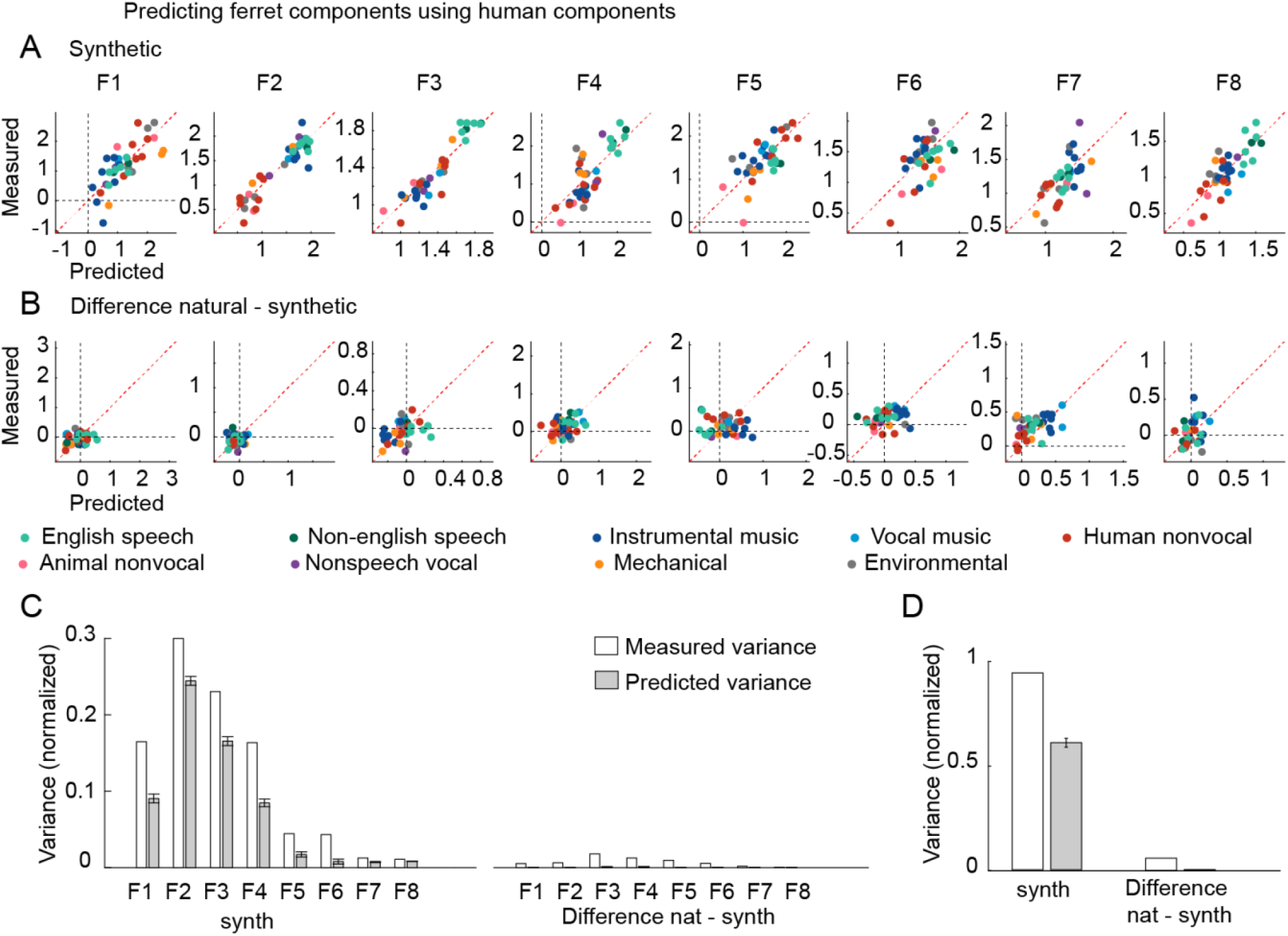
Results of predicting ferret components from human components. Same format as **Fig S6B-E**.

**Figure S8.**
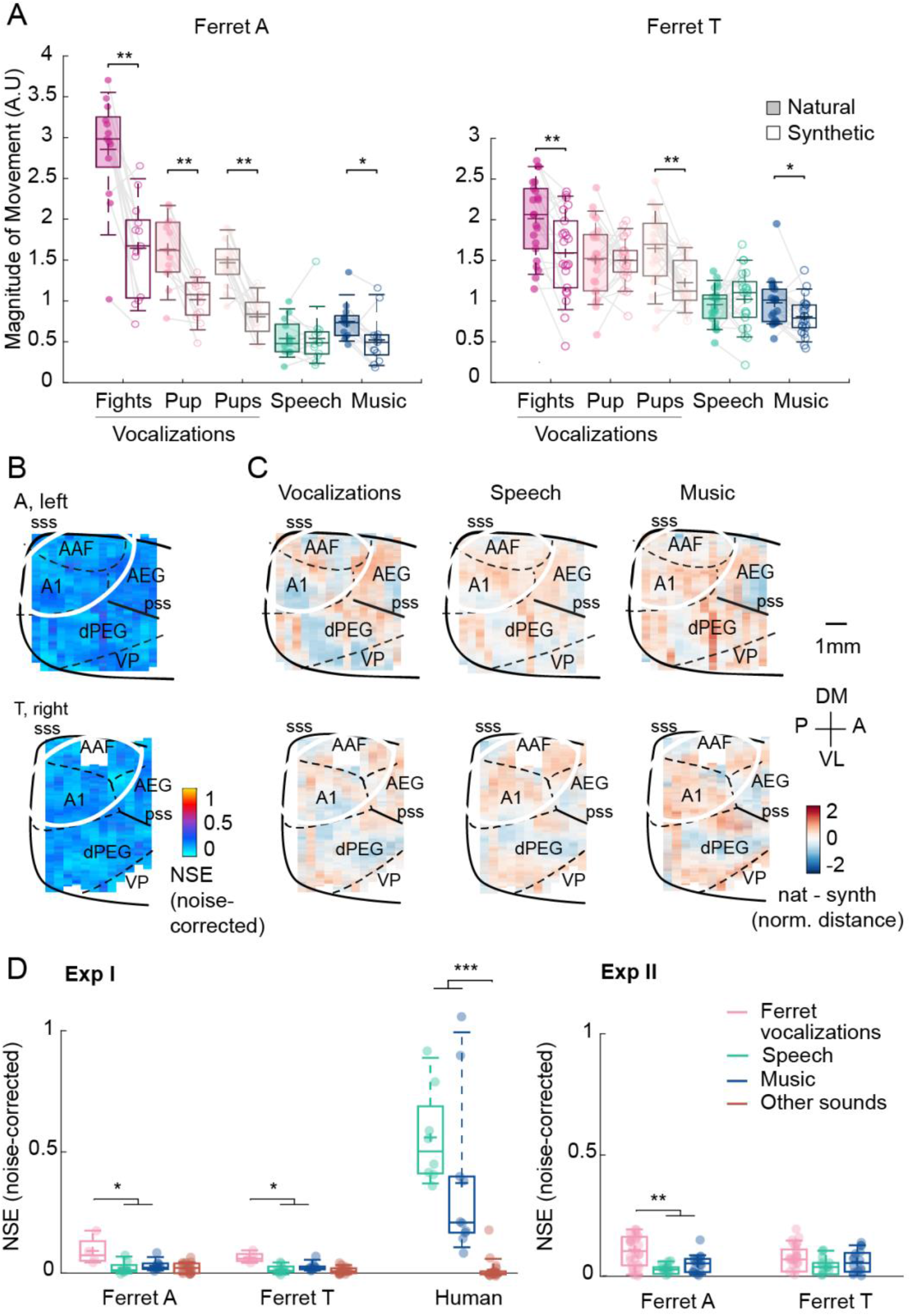
Results of Experiment II from other hemispheres. **A-C**, Same format as **Fig 4C-E**, except that in panel A the vocalizations are split into sub-categories: fight calls, single pup calls, multiple pup calls. Movement amplitude is shown for each animal separately. **D**, This panel shows the distribution of NSE values for all pairs of natural and synthetic sounds (median across all voxels), grouped by category. The numerator in the NSE calculation is simply the squared error for that sound pair, and the denominator is computed in the normal way using responses to all sounds (equation 1). Dots show individual sound pairs and box-plots show the median, central 50% and central 92% (whiskers) of the distribution.

**Figure S9.**
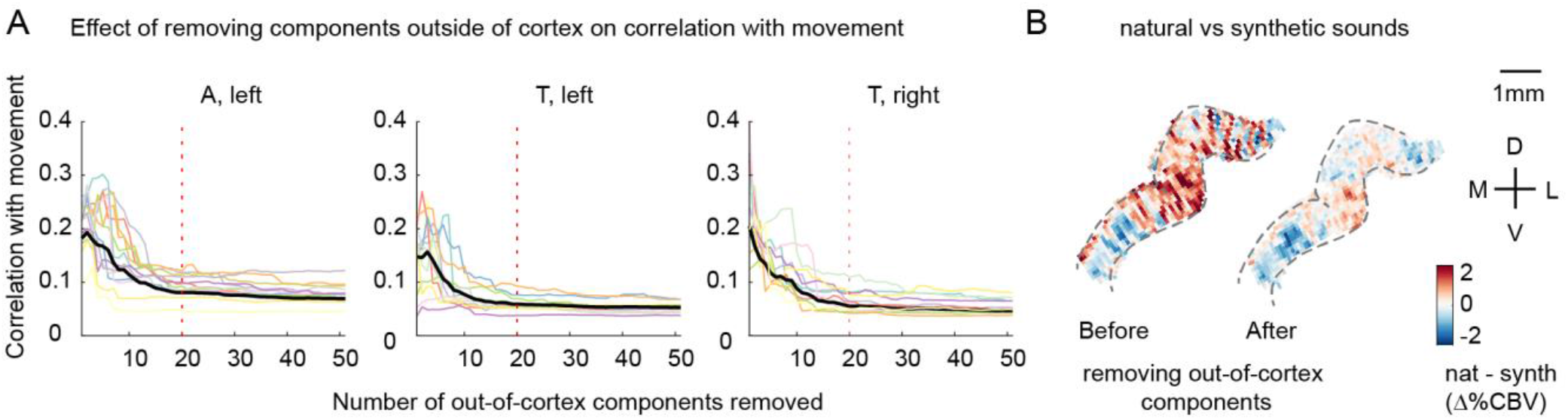
The effect of removing outside-of-cortex components on motion correlations. Voxel responses were denoised by removing components from outside of cortex, which are likely to reflect artifacts like motion (see Denoising Part I in Methods). **A**, Effect of removing components from outside of cortex on correlations with movement. We measured the correlation of each voxel’s response with movement, measured from a video recording of the animal’s face (absolute deviation between adjacent frames). Each line shows the average absolute correlation across voxels for a single recording session / slice. Correlation values are plotted as a function of the number of removed components. Motion correlations were substantially reduced by removing the top 20 components (vertical dotted line). **B**, The average difference between responses to natural vs synthetic sounds for an example slice before and after removing the top 20 out-of-cortex components. Motion induces a stereotyped “striping” pattern due to its effect on blood vessels, which is evident in the map computed from raw data, likely because ferrets moved substantially more during natural vs. synthetic sounds (particular for ferret vocalizations; **Figure 4C**). The striping pattern is largely removed by the denoising procedure.

## Appendix Recentered CCA

### Derivation

The goal of the denoising procedure described in Part I was to remove artifactual components that were present both inside and outside of cortex, since such components are both likely to be artifactual and likely to distort the responses-of-interest. The key complication was that motion-induced artifacts are likely to be correlated with true sound-driven neural activity because the animals reliably moved more during the presentation of some sounds. To deal with this issue, we used the fact that motion will vary from trial-to-trial for repeated presentations of the same sound, while sound-driven responses by definition will not. Here, we give a more formal derivation of our procedure. We refer to our method as “recentered CCA” (rCCA) for reasons that will become clear below.

We represent the data for each voxel as an unrolled vector (***d***_***v***_) that contains its response timecourse across all sounds and repetitions. We assume these voxel responses are contaminated by a set of K artifactual component timecourses {***a***_***k***_}. We thus model each voxel as a weighted sum of these artifactual components plus a sound-driven response timecourse (***s***_***v***_):

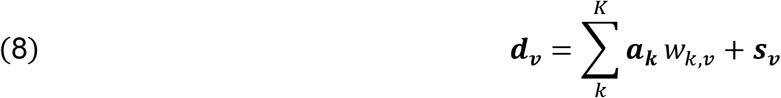

Actual voxel responses are also corrupted by voxel-specific noise, which would add an additional error term to the above equation. In practice, the error term has no effect on our derivation so we omit it for simplicity (we verified our analysis was robust to voxel-specific noise using simulations, which are described below).

To denoise our data, we need to estimate the artifactual timecourses {***a***_***k***_} and their weights (*w*_*k,v*_) so that we can subtract them out. If the artifactual components {***a***_***k***_} were uncorrelated with the sound-driven responses (***s***_***v***_) we could estimate them by performing CCA on voxel responses from inside and outside of cortex, since only the artifacts would be correlated. However, we expect sound-driven responses to be correlated with motion artifacts, and the components inferred by CCA will thus reflect a mixture of sound-driven and artifactual activity.

To overcome this problem, we first subtract-out the average response of each voxel across repeated presentations of the same sound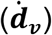. This “recentering” operation removes sound-driven activity, which by definition is the same across repeated presentations of the same sound:

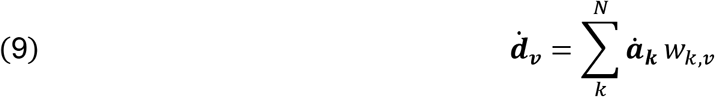

where the dot above a variable indicates its response after recentering (not its time derivative). Because sound-driven responses have been eliminated, applying CCA to the recentered voxel responses should yield an estimate of the recentered artifacts 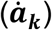 and their weights (*w*_*k,v*_) (note that CCA actually yields a set of components that span a similar subspace as the artifactual components, which is equivalent from the perspective of denoising). To simplify notation in the equations below, we assume this estimate is exact (i.e. CCA exactly returns 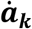 and *w*_*k,v*_).

Since the weights (*w*_*k,j*_) are the same for original (***d***_***v***_) and recentered 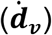 data, we are halfway done. All that is left is to estimate the original artifact components before recentering (***a***_***k***_), which can be done using the original data before recentering (***d***_***v***_)). o see this, first note that canonical components are by construction a linear projection of the data used to compute them, and thus, we can write:

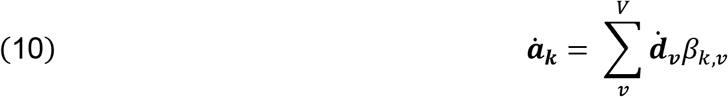

We can use the reconstruction weights (*β*_*k,v*_) in the above equation to get an estimate of the original artifactual components by applying them to the original data before recentering:

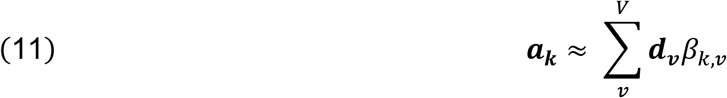

To see this, we expand the above equation:

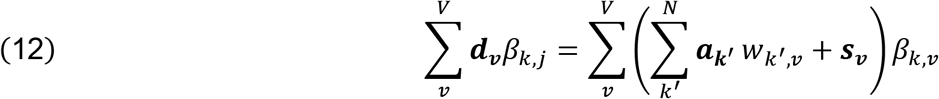

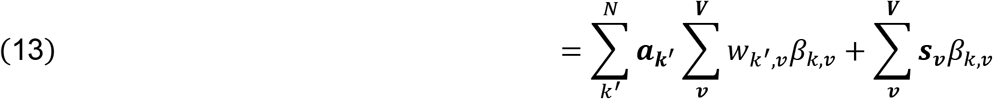

The first term in the above equation exactly equals ***a***_***k***_ because *wk*_*′*_,*v* and *β*_*k,v*_ are by construction pseudoinverses of each other (i.e. 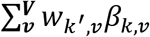 is 1 when *k*′ = *k* and 0 otherwise). The second term can be made small by estimating and applying reconstruction weights using only data from outside of cortex, where sound-driven responses are weak.

We thus have a procedure for estimating both the original artifactual responses (***a***_***k***_) and their weights (*w*_*k,j*_), and can denoise our data by simply subtracting them out:

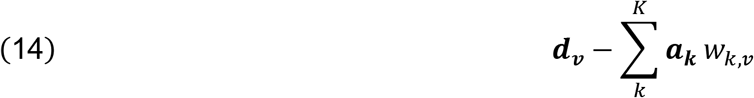

### Procedure

We now give the specific steps used to implement the above procedure using matrix notation. The inputs to the analysis were two matrices (*D*_*in*_, *D*_*out*_), each of which contained voxel responses from inside and outside of cortex. Each column of each matrix contained the response timecourse of a single voxel, concatenated across all sounds and repetitions (i.e. ***d***_***v***_ in the above derivation). We also computed recentered data matrices 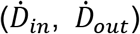 by subtracting out trial-averaged activity (i.e. 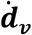).

CCA can be performed by whitening each input matrix individually, concatenating the whitened data matrices, and then computing the principal components of the concatenated matrices (de Cheveigné et al., 2019). Our procedure is an elaborated version of this basic design:

1. The recentered data matrices were reduced in dimensionality and whitened. We implemented this step using the singular value decomposition (SVD), which factors the data matrix as the product of two orthonormal matrices (*U* and *V*), scaled by a diagonal matrix of singular values (*S*):

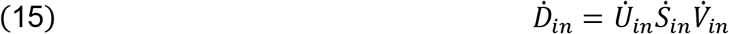

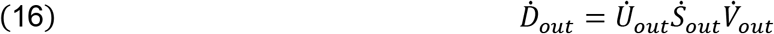

The reduced and whitened data was given by selecting the top 250 components and removing the diagonal S matrix:

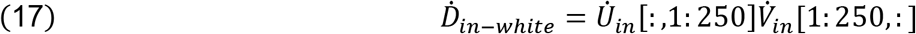

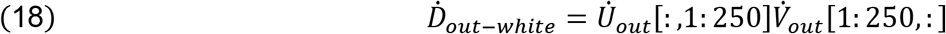
2. We concatenated the whitened data matrices from inside and outside of cortex across the voxel dimension:

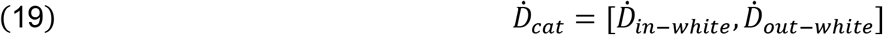
3. We computed the top N principal components from the concatenated matrix using the SVD:

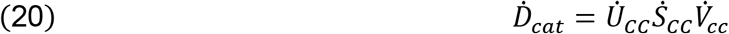 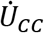 contains the timecourses of the canonical components (CCs), ordered by variance, which provide an estimate of the artifactual components after recentering (i.e. 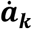). The corresponding weights (i.e. *w*_*k,v*_) for voxels inside of cortex were computed by projecting the recentered data onto 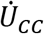:

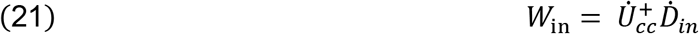

where + indicates the matrix pseudo-inverse.
4. The original artifactual components before recentering (i.e. ***a***_***k***_) were estimated by learning a set of reconstruction weights (B) using recentered data from outside of cortex, and then applying these weights to the original data before recentering:

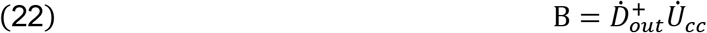

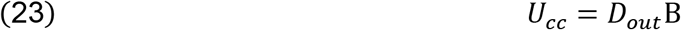

*U*_*cc*_ is an estimate of the artifactual components before recentering (i.e. ***a***_***k***_).
5. Finally, we subtracted out the contribution of the artifactual components to each voxel inside of cortex, estimated by simply multiplying the component responses and weights:

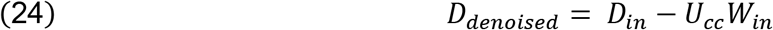

### Simulation

We created a simple simulation to test our method. We simulated 1000 voxel responses, both inside and outside of cortex, using equation 8. For voxels outside of cortex, we set the sound-driven responses to 0. We also added voxel-specific noise to make the denoising task more realistic/difficult (sampled from a Gaussian). Results were very similar across a variety of noise levels.

To induce correlations between the artifactual (***a***_***k***_) and sound-driven responses (***s***_***v***_), we forced them to share a subspace. Specifically, we computed the sound-driven responses as a weighted sum of a set of 10 component timecourses (results did not depend on this parameter), thus forcing the responses to be low-dimensional, as we found to be the case:

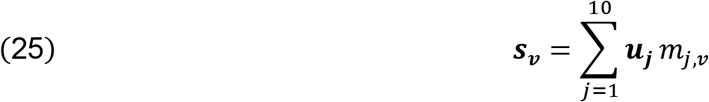

The artifactual timecourses were then computed as a weighted sum of these same 10 components timecourses plus a timecourse that was unique to each artifactual component:

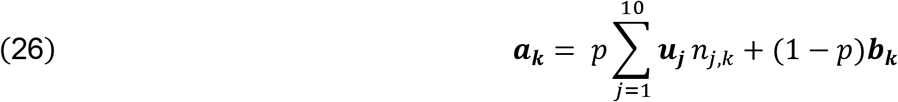

where *p* controls the strength of the dependence between the sound-driven and artifactual components with a value of 1 indicating complete dependence and 0 indicating no dependence. All of responses and weights (***u***_***j***_, ***b***_***k***_, *m*_*j,v*_, *n*_*j,k*_) were sampled from a unit-variance Gaussian. Sound-driven responses were constrained to be the same across repetitions by sampling the latent timecourses ***u***_***j***_ once per sound, and then simply repeating the sampled values across repetitions. In contrast, a unique ***b***_***k***_ was sampled for every repetition of every sound to account for the fact that the artifacts like motion will vary from trial-to-trial. We sampled 20 artifactual timecourses using equation 26.

We applied both standard CCA and our modified rCCA method to the simulated data. We measured the median NSE between the true and estimated sound-driven responses (***s***_***v***_), computed using the two methods as a function of the strength of the dependence (*p*) between sound-driven and artifactual timecourses (**Fig A1A**). For comparison, we also plot the NSE for raw voxels (i.e. before any denoising) as well as the minimum possible NSE (noise floor) given the voxel-specific noise (which cannot possibly be removed using CCA or rCCA). When the dependence is low, both CCA and rCCA yield similarly good results, as expected. As the dependence increases, CCA performs substantially worse, while rCCA continues to perform well up until the point when the dependence becomes so strong that sound-driven and artifactual timecourses are nearly indistinguishable. Results were not highly sensitive to the number of components removed as long as the number of removed components was equal to or greater than the number of artifactual components (**Figure A1B**).

**Figure A1:**
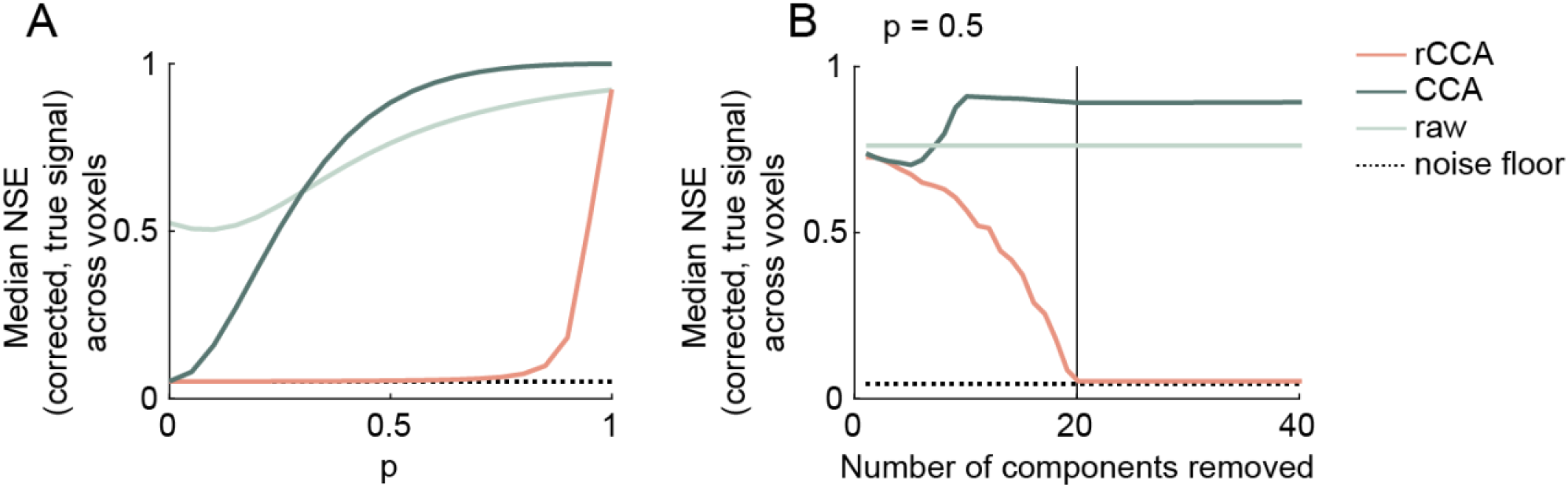
Simulation results. **A**. Median NSE across simulated voxels between the true and estimated sound-driven responses (***s***_***v***_), computed using raw/undenoised data (light green line), standard CCA (dark green line), and recentered CCA (red line). Results are shown as a function of the strength of the dependence (*p*) between sound-driven and artifactual timecourses. The minimum possible NSE (noise floor) given the level of voxel-specific noise is also shown. **B**. Same as panel A, but showing results as a function of the number of components removed for a fixed value of *p* (set to 0.5).

## References

Agamaite JA, Chang C-J, Osmanski MS, Wang X (2015) A quantitative acoustic analysis of the vocal repertoire of the common marmoset (Callithrix jacchus). The Journal of the Acoustical Society of America 138:2906–2928.

Belin P, Zatorre RJ, Lafaille P, Ahad P, Pike B (2000) Voice-selective areas in human auditory cortex. Nature 403:309–312.

Bimbard C, Demene C, Girard C, Radtke-Schuller S, Shamma S, Tanter M, Boubenec Y (2018) Multi-scale mapping along the auditory hierarchy using high-resolution functional UltraSound in the awake ferret. Elife 7:e35028.

Boebinger D, Norman-Haignere S, McDermott J, Kanwisher N (2020) Cortical music selectivity does not require musical training. bioRxiv.

Brodbeck C, Hong LE, Simon JZ (2018) Rapid transformation from auditory to linguistic representations of continuous speech. Current Biology 28:3976–3983.

Bruns V, Schmieszek E (1980) Cochlear innervation in the greater horseshoe bat: demonstration of an acoustic fovea. Hearing research 3:27–43.

Chi T, Ru P, Shamma SA (2005) Multiresolution spectrotemporal analysis of complex sounds. The Journal of the Acoustical Society of America 118:887–906.

de Cheveigné A, Di Liberto GM, Arzounian D, Wong DD, Hjortkjær J, Fuglsang S, Parra LC (2019) Multiway canonical correlation analysis of brain data. NeuroImage 186:728–740.

de Cheveigné A, Parra LC (2014) Joint decorrelation, a versatile tool for multichannel data analysis. Neuroimage 98:487–505.

de Heer WA, Huth AG, Griffiths TL, Gallant JL, Theunissen FE (2017) The hierarchical cortical organization of human speech processing. Journal of Neuroscience:3267–16.

Demené C, Deffieux T, Pernot M, Osmanski B-F, Biran V, Gennisson J-L, Sieu L-A, Bergel A, Franqui S, Correas J-M (2015) Spatiotemporal clutter filtering of ultrafast ultrasound data highly increases Doppler and fUltrasound sensitivity. IEEE transactions on medical imaging 34:2271–2285.

Di Liberto GM, Wong D, Melnik GA, de Cheveigné A (2019) Low-frequency cortical responses to natural speech reflect probabilistic phonotactics. Neuroimage 196:237–247.

DiCarlo JJ, Cox DD (2007) Untangling invariant object recognition. Trends in cognitive sciences 11:333–341.

Ding N, Patel AD, Chen L, Butler H, Luo C, Poeppel D (2017) Temporal modulations in speech and music. Neuroscience & Biobehavioral Reviews.

Eliades SJ, Miller CT (2017) Marmoset vocal communication: behavior and neurobiology. Developmental neurobiology 77:286–299.

Erb J, Armendariz M, De Martino F, Goebel R, Vanduffel W, Formisano E (2019) Homology and specificity of natural sound-encoding in human and monkey auditory cortex. Cerebral Cortex 29:3636–3650.

Gesnik M, Blaize K, Deffieux T, Gennisson J-L, Sahel J-A, Fink M, Picaud S, Tanter M (2017) 3D functional ultrasound imaging of the cerebral visual system in rodents. NeuroImage 149:267–274.

Hickok G, Poeppel D (2007) The cortical organization of speech processing. Nature reviews neuroscience 8:393–402.

Joris PX, Bergevin C, Kalluri R, Mc Laughlin M, Michelet P, van der Heijden M, Shera CA (2011) Frequency selectivity in Old-World monkeys corroborates sharp cochlear tuning in humans. Proceedings of the National Academy of Sciences 108:17516–17520.

Kell AJ, Yamins DL, Shook EN, Norman-Haignere SV, McDermott JH (2018) A task-optimized neural network replicates human auditory behavior, predicts brain responses, and reveals a cortical processing hierarchy. Neuron.

Köppl C, Gleich O, Manley GA (1993) An auditory fovea in the barn owl cochlea. Journal of Comparative Physiology A 171:695–704.

Leonard MK, Bouchard KE, Tang C, Chang EF (2015) Dynamic encoding of speech sequence probability in human temporal cortex. Journal of Neuroscience 35:7203–7214.

Macé E, Montaldo G, Cohen I, Baulac M, Fink M, Tanter M (2011) Functional ultrasound imaging of the brain. Nature methods 8:662.

McDermott JH, Simoncelli EP (2011) Sound texture perception via statistics of the auditory periphery: evidence from sound synthesis. Neuron 71:926–940.

Mesgarani N, Cheung C, Johnson K, Chang EF (2014) Phonetic feature encoding in human superior temporal gyrus. Science 343:1006–1010.

Mesgarani N, David SV, Fritz JB, Shamma SA (2008) Phoneme representation and classification in primary auditory cortex. The Journal of the Acoustical Society of America 123:899–909.

Milham MP, Ai L, Koo B, Xu T, Amiez C, Balezeau F, Baxter MG, Blezer EL, Brochier T, Chen A (2018) An open resource for non-human primate imaging. Neuron 100:61–74.

Mizrahi A, Shalev A, Nelken I (2014) Single neuron and population coding of natural sounds in auditory cortex. Current opinion in neurobiology 24:103–110.

Moore JM, Woolley SM (2019) Emergent tuning for learned vocalizations in auditory cortex. Nature neuroscience 22:1469–1476.

Nelken I, Bizley JK, Nodal FR, Ahmed B, King AJ, Schnupp JW (2008) Responses of auditory cortex to complex stimuli: functional organization revealed using intrinsic optical signals. Journal of neurophysiology 99:1928–1941.

Norman-Haignere SV, Kanwisher N, McDermott JH, Conway BR (2019) Divergence in the functional organization of human and macaque auditory cortex revealed by fMRI responses to harmonic tones. Nature Neuroscience 22:1057.

Norman-Haignere SV, Kanwisher NG, McDermott JH (2015) Distinct cortical pathways for music and speech revealed by hypothesis-free voxel decomposition. Neuron 88:1281–1296.

Norman-Haignere SV, McDermott JH (2018) Neural responses to natural and model-matched stimuli reveal distinct computations in primary and nonprimary auditory cortex. PLoS biology 16:e2005127.

Overath T, McDermott JH, Zarate JM, Poeppel D (2015) The cortical analysis of speech-specific temporal structure revealed by responses to sound quilts. Nature neuroscience 18:903–911.

Patel AD (2012) Language, music, and the brain: a resource-sharing framework. Language and music as cognitive systems:204–223.

Petkov CI, Kayser C, Steudel T, Whittingstall K, Augath M, Logothetis NK (2008) A voice region in the monkey brain. Nature neuroscience 11:367–374.

Polley DB, Steinberg EE, Merzenich MM (2006) Perceptual Learning Directs Auditory Cortical Map Reorganization through Top-Down Influences. J Neurosci 26:4970–4982.

Schnupp JW, Hall TM, Kokelaar RF, Ahmed B (2006) Plasticity of temporal pattern codes for vocalization stimuli in primary auditory cortex. Journal of Neuroscience 26:4785–4795.

Singh NC, Theunissen FE (2003) Modulation spectra of natural sounds and ethological theories of auditory processing. The Journal of the Acoustical Society of America 114:3394–3411.

Srihasam K, Vincent JL, Livingstone MS (2014) Novel domain formation reveals proto-architecture in inferotemporal cortex. Nature neuroscience 17:1776.

Steinschneider M, Nourski KV, Fishman YI (2013) Representation of speech in human auditory cortex: is it special? Hearing research 305:57–73.

Theunissen FE, Elie JE (2014) Neural processing of natural sounds. Nature Reviews Neuroscience 15:355–366.

Walker KM, Gonzalez R, Kang JZ, McDermott JH, King AJ (2019) Across-species differences in pitch perception are consistent with differences in cochlear filtering. eLife 8:e41626.

Zatorre RJ, Belin P, Penhune VB (2002) Structure and function of auditory cortex: music and speech. Trends in Cognitive Sciences 6:37–46.

